# Angiopoietin signalling is a central axis of amyloid-driven vascular dysfunction in Alzheimer’s disease

**DOI:** 10.1101/2025.08.25.672093

**Authors:** Matthias Flotho, Andrew Yang, Fabian Kern, Simon Graf, Ian Ferenc Diks, Heather Shin, Kristy A. Zera, Daniela Berdnik, Maayan R. Agam, Divya Channappa, Sophia M. Shi, Malia Alexandra Belnap, Epiphani C. Simmons, Karen Bradshaw, Friederike Grandke, David A. Bennett, Marion Buckwalter, Andreas Keller, Tony Wyss-Coray

## Abstract

The neurovascular unit is critical for brain health, and its dysfunction has been linked to Alzheimer’s disease (AD). However, a cell-type-resolved understanding of how diverse vascular cells become dysfunctional and contribute to disease has been missing. Here, we applied Vessel Isolation and Nuclei Extraction for Sequencing (VINE-seq) to build a comprehensive transcriptomic atlas from 101 individuals along AD progression. Our analysis of over 842,646 parenchymal and vascular nuclei reveals that vascular dysfunction in AD is driven by transcriptional changes rather than shifts in cell proportions, with brain endothelial cells (BECs) and smooth muscle cells (SMCs) most affected. Strikingly, these molecular signatures emerge early at the mild cognitive impairment (MCI) stage, implicating vascular dysfunction early in AD pathogenesis. Stratifying by pathology reveals distinct vascular responses to β-amyloid and tau: β-amyloid burden primarily perturbs BECs and SMCs, while tau pathology predominantly impacts glial cells. We identify dysregulated angiopoietin signaling across multiple vascular cell types as a key axis, with antagonistic ANGPT2 in vascular cells and ANGPT1 in astrocytes becoming progressively dysregulated with AD. Together, this work provides a foundational resource that reveals early and pathology-specific pathways of vascular dysfunction in AD.

**Key Messages:**

- VINE-seq analysis from 101 individuals creates a comprehensive human brain vascular atlas across Alzheimer’s disease (AD) progression.
- AD vascular dysfunction is driven by transcriptional changes rather than shifts in cell proportions, with BECs and SMCs most affected.
- Transcriptional signatures of vascular dysfunction emerge early at the mild cognitive impairment (MCI) stage, preceding severe cognitive symptoms and aligning more closely with AD than cognitively normal individuals.
- Aβ and tau associate with distinct vascular changes: Aβ mainly perturbs endothelial and smooth muscle cells, while tau impacts microglia and astrocytes.
- Angiopoietin signaling (antagonistic ANGPT2 in vascular cells vs. ANGPT1 in astrocytes) becomes progressively dysregulated during AD progression.

## INTRODUCTION

Proper function of the brain’s vasculature is essential for supporting brain homeostasis and overall neurological health^1–4^. Dysfunction of this intricate cellular network is a key feature of brain aging and a major contributor to various neurodegenerative disorders, including Alzheimer’s disease (AD)^5–12^. However, the molecular mechanisms that underlie vascular contributions to AD remain poorly understood.

In recent years, large-scale single-cell RNA sequencing (scRNA-seq) studies of the human brain have revolutionized our understanding of AD, revealing cell-type-specific gene expression changes that underlie disease progression^13–21^. Major consortia and reference cohorts, such as the Religious Orders Study and Rush Memory and Aging Project (ROSMAP)^8^ and the Seattle Alzheimer’s Disease Brain Cell Atlas (SEA-AD), have generated vast datasets comprising millions of cells. These resources have provided unprecedented insights into the biology of neural and glial cells in AD.

Despite this progress, the cellular and molecular pathways of neurovascular dysfunction remain poorly characterized along AD progression. Most large-scale brain atlases have focused on parenchymal cells^13–24^, neurons and glia, leaving the diverse populations of endothelial cells, pericytes, smooth muscle cells, and associated immune cells that constitute the vasculature largely underrepresented^25–27^. This represents a profound data gap, as the health of these vascular cells is paramount for regulating cerebral blood flow, maintaining the integrity of the blood-brain barrier (BBB), and clearing metabolic waste, including amyloid-β^1–4,28–30^. Understanding how these fundamental vascular functions are altered at the molecular level in AD is therefore a critical unmet need.

To fill this gap, we applied our Vessel Isolation and Nuclei Extraction for Sequencing (VINE-Seq) protocol^31^ to generate a deep transcriptomic atlas of the neurovascular unit from 101 individuals in ROSMAP spanning the full clinical and pathological spectrum from normality to AD. This high-resolution resource allowed us to systematically investigate the molecular drivers of vascular dysfunction at the single-cell level. Specifically, we dissected the cell-type-specific transcriptional responses to distinct neuropathological lesions (β-amyloid and tau) and compared these signatures with those associated with clinical stages of cognitive decline^32,33^. Our findings reveal early and pathology-specific dysregulation of vascular programs, identifying the angiopoietin signaling axis as a central mediator of neurovascular dysfunction in AD.

## RESULTS

### A deep transcriptomic map of brain vascular cells exposed to AD pathology

Towards understanding how human blood vessels change at the molecular level in the context of AD pathology we used our previously published method, VINE-Seq^31^, to process 110 brains of individuals from the ROSMAP cohort^34^ with different degrees of pathology and clinical diagnosis (**Supplemental Table 1**). We sequenced RNA isolated from nuclei associated with vascular cells from the frontal cortex of samples and after excluding brains with low yields of vascular cells **(Methods)** ended up with 101 brains from individuals with different degrees of AD pathology and clinical diagnosis of NCI (n = 39), MCI (n = 31), and AD (n = 31) **(*Fig. 1a*).** Brains were equally distributed between men and women and postmortem intervals were similar (***Supplemental Fig. 1a,b)***. After quality control we obtained 842,646 single nucleus transcriptomes which were separated based on established marker genes into ten broadly defined vascular and parenchymal cell types and visualized using uniform manifold approximation and projection (UMAP) space **(*Fig. 1b* and *Supplemental Fig. 2)***. A more detailed annotation yielded 435,845 high-quality vascular cell nuclei allowing us to describe molecular changes in the human brain vasculature at unprecedented depth **(Fig. 1c,d and *Supplemental Fig. 3)***. We thus obtained arterial, capillary, and venous brain endothelial cells (BECs), arterial and arteriolar smooth muscle cells (SMCs), meningeal and perivascular fibroblast, and our recently described M and T pericytes^31^ at numbers far exceeding those of previous studies. Additionally, we identified distinct immune cell type clusters including perivascular macrophages, microglia, and T cells. Generally, cell type proportions were similar across neuropathological or clinical stages of disease and age of death **(*Supplemental Fig. 4a*)** and while we observed fewer BECs with increasing β-amyloid load and more reactive microglia with higher neuritic plaque counts ***(Supplemental Fig. 4b,c)***, these numbers were not significant after adjustment for multiple hypothesis testing across cell types and neuropathological or clinical variables.

**Fig. 1.**
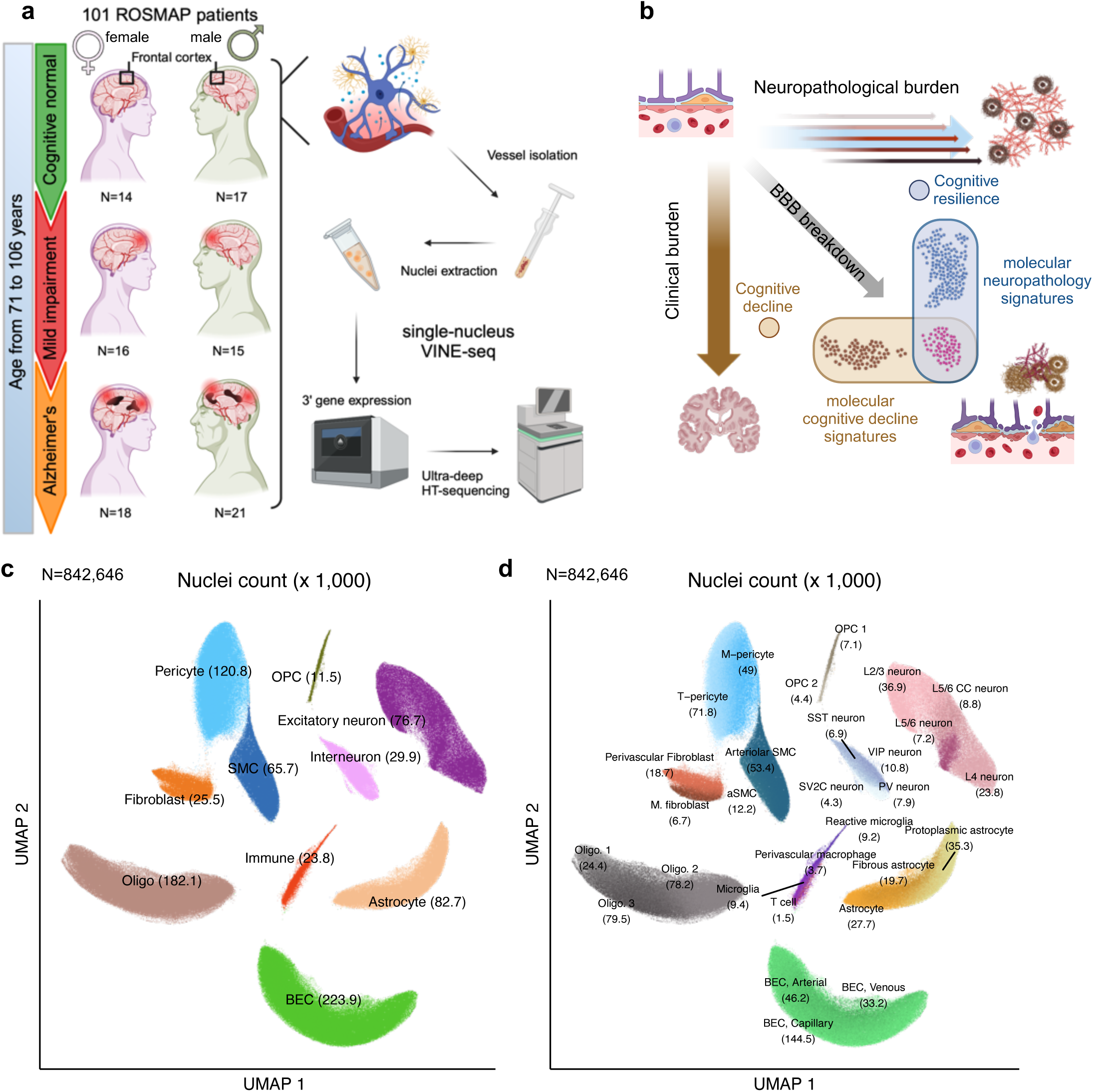
Cohort description and cell-type composition: **(a)** Schematic view of the cohort (left) and sample processing using the VINE-seq (Vessel Isolation and Nuclei Extraction for Sequencing) protocol with droplet-based sequencing (right). Samples are balanced with respect to sex (female: purple; male: green) and the different clinical diagnoses. **(b)** Representation of the relationship between neuropathological burden (horizontal direction; small arrows indicate different neuropathological markers such as PHFtau tangle density or β-amyloid load, thick arrow indicates the combined NIA-Reagan pathologic AD), clinical diagnosis, and cognitive outcomes in Alzheimer’s disease (vertical direction), illustrating the connections and potential overlaps (diagonal arrow).**(c)** UMAP (Uniform Manifold Approximation and Projection) of the 842,646 high-quality nuclei in the dataset, with broad cell type labeling (10 cell types). Vasculature cells are highlighted in green (BEC), light blue (Pericytes), and dark blue (SMCs). **(d)** Same UMAP as in **(c)** but with fine cell type labeling (29 cell types identified in total).

To determine transcriptional differences of vascular cells associated broadly with AD neuropathology we used the NIA-Reagan diagnosis to separate participants into those with and without pathologic AD^12^, which is ascertained by the presence and distribution of neurofibrillary tangles and neuritic plaques^35^ following standard procedures^36,37^ ***(Supplemental Fig. 5)***. Across 15 cell types assessed, meningeal fibroblasts showed twice as many differentially expressed genes (DEGs; adjusted p<0.05) than the next cell type, reactive microglia **(Fig. 2a and *Supplemental Table 2*)** with most cell types showing fewer than 30 DEGs. These small numbers of DEG counts are likely the result of restrictive parameter selections in the MAST pipeline, setting the defaults to logFC>0.25 and the confounder to sex. The genes most significantly different between brain with and without pathologic AD are predominantly expressed in BECs with ADAMTS9, PDXK, STEAP1B, ARL17B, ZFY, and ANGPT2 overlapping among arterial, capillary, and venous BECs **(Fig. 2b,c)**. Among these differentially expressed genes in BECs, ANGPT2 stood out as the strongest upregulated gene in capillary BECs (logFC=0.59; p<10^-200^). Compared to other cell types, the BEC subtypes exhibited DEGs with the highest significance, partially due to the high number of cells captured in our dataset **(*Fig. 2b*)**. Beyond BECs, other notable dysregulated genes included *PLCG2*, which has been linked to AD and longevity^38,39^, broadly upregulated across pericytes, astrocytes, SMCs, and perivascular fibroblasts.

**Fig. 2.**
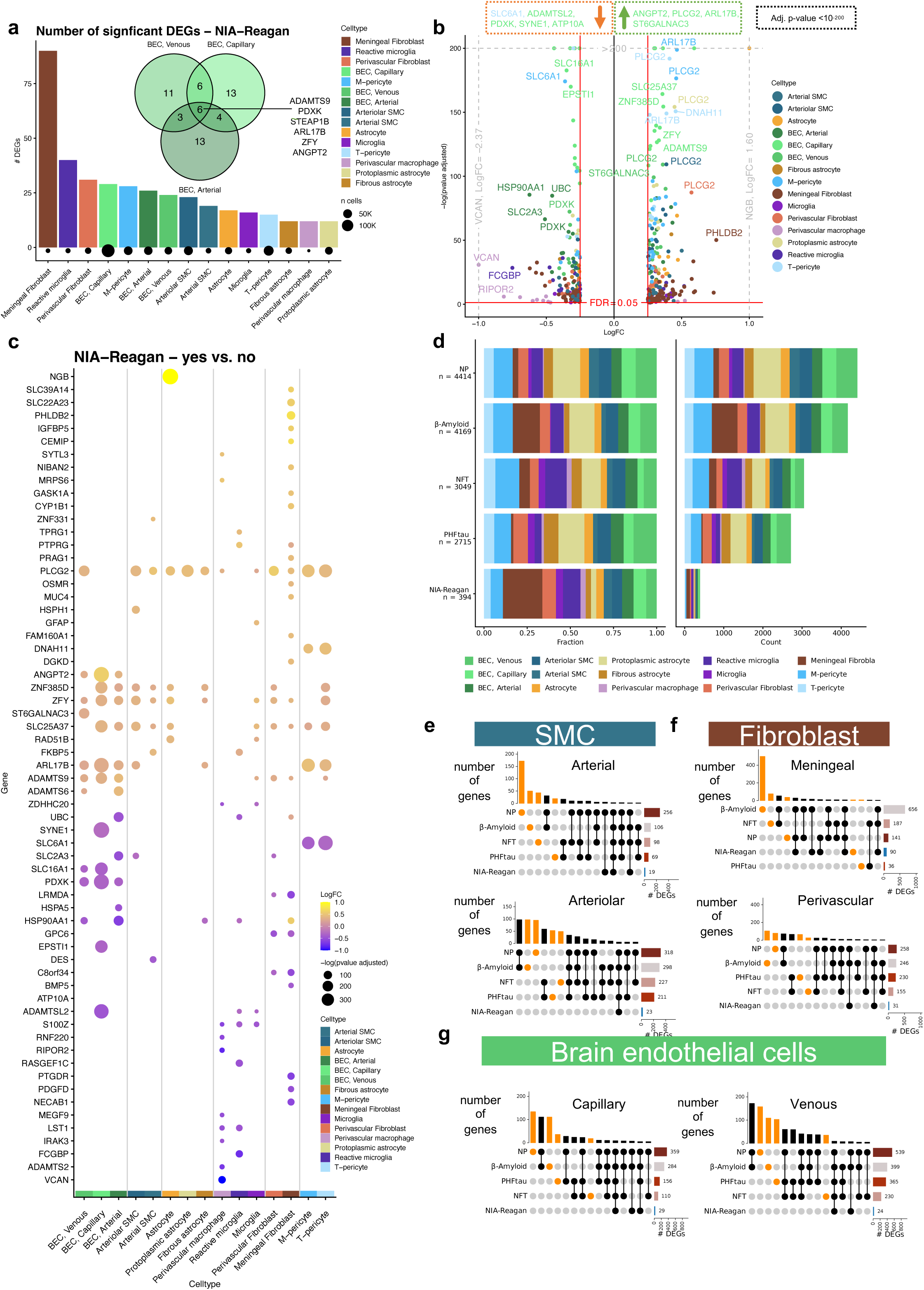
Gene signatures related to neuropathology in vascular cell types: **(a)** Bar plot showing the number of significant differentially expressed genes (DEGs, FDR < 0.05) between NIA-Reagan AD diagnosed and healthy persons in vascular cell types. The Venn diagram highlights the DEG overlaps within the BEC sub-cell types. **(b)** A volcano plot (x-axis as log fold change, y-axis as negative decadic logarithm of the p-value) showing all significant genes colored with respect to the cell type of origin. **(c)** Dot plot displaying the DEGs grouped by fine-cell type, showing similar patterns within the broader cell type definition. The dot size represents the significance (negative decadic logarithm of the p-value), and the color the log fold change. **(d)** Scaled and absolute numbers of DEGs for the different neuropathological lesions and diagnoses. For the lesion-specific DEGs, the lowest third of lesion abundance was compared with the third with the highest abundance. **(e)** Upset plots highlighting the DEGs between the different neuropathological burdens in arterial and arteriolar SMCs. Orange bars denote gene sets that are unique to one neuropathological marker. **(f)** Analogous to **(e)**, upset plot for the fibroblast cell types. **(g)** Analogous to (**e**), upset plots for capillary and venous BEC sub-cell types.

The modest number of differentially expressed genes (DEGs) associated with the composite NIA-Reagan score suggested that this broad categorization may obscure molecular associations with specific lesion types. We therefore performed analyses comparing individuals in the highest versus lowest tertiles for three distinct neuropathological measures: β-amyloid load, neuritic plaque (NP) counts, and neurofibrillary tangle (NFT) density (***Supplemental Fig. 6a***)^8,11,12,40–42^.

This granular approach revealed an order-of-magnitude more DEGs than the binary NIA-Reagan comparison, with the strongest associations linked to amyloid-related pathologies. Specifically, analyses of β-amyloid load and NP counts identified 4,169 and 4,414 DEGs, respectively, whereas the analysis of NFT density yielded substantially fewer (***Fig. 2d, Supplemental Table 3***). These findings indicate that in the cell types of the neurovascular unit, transcriptional responses are more strongly associated with the presence of amyloid pathologies than with tau pathology.

The DEGs associated with β-amyloid load and NP counts were overwhelmingly concentrated in vascular cells—namely, brain endothelial cells (BECs), smooth muscle cells (SMCs), and fibroblasts. This analysis highlighted specific cellular associations: meningeal fibroblasts, for example, showed a strong transcriptional response specifically associated with high β-amyloid load (***Fig. 2f***), while arterial SMCs had a high number of DEGs uniquely associated with NP counts (***Fig. 2e***). Across all BEC subtypes, the largest number of DEGs was associated with high NP counts and high β-amyloid load (***Fig. 2g***). Across these lesion-specific analyses, *ANGPT2* consistently emerged as a top upregulated gene in BECs (***Supplemental Fig. 7***). Its antagonist, *ANGPT1*,^43^ was not broadly dysregulated in these comparisons, informing a more targeted investigation into the angiopoietin axis.

### ANGPTs in SMC and BEC contribute to AD neuropathology

Given that ANGPT2 was a top differentially expressed gene (DEG) associated with amyloid pathology, we performed a targeted analysis of the angiopoietin signaling axis across different neuropathological measures. Examination of gene expression across tertiles of neuropathological burden revealed distinct, opposing patterns for *ANGPT2* and its antagonist *ANGPT1*^43^. In vascular cells like arterial SMCs, *ANGPT2* expression was higher in individuals in both the medium and high burden groups compared to the low burden group (***Fig. 3a***). In contrast, *ANGPT1* was most pronounced in astrocytes only in the group with the highest pathological burden (***Fig. 3b***).

**Fig. 3.**
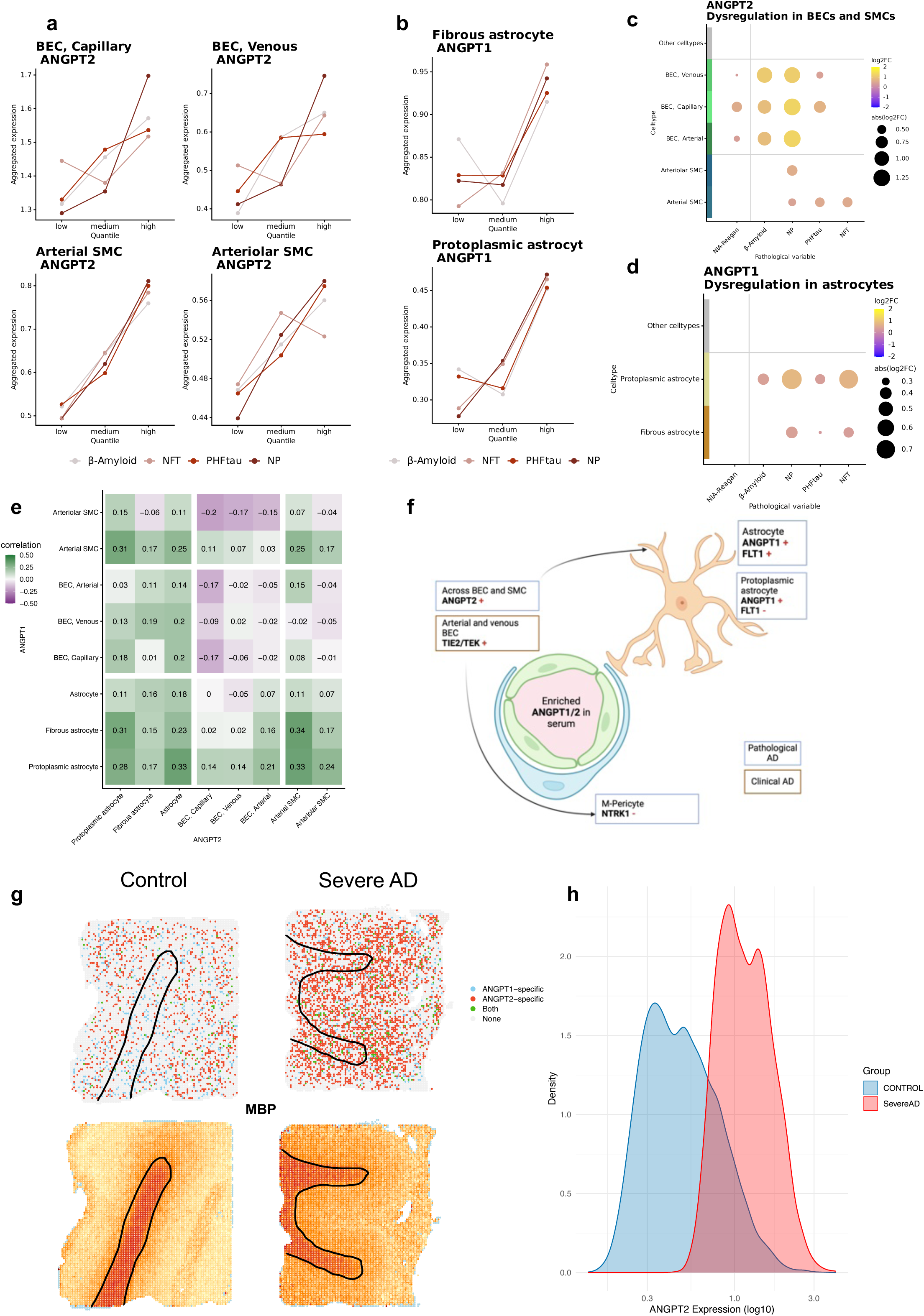
Signs of BBB-breakdown in high-resolution pathology variables: **(a)** Pseudobulk gene expression across the person’s lesion progression (represented by distinct colors). The cohort is split into three quantiles for each neuropathological lesion. For ANGPT2, selected BEC and SMC sub-types are displayed. **(b)** Analogous to (a), pseudobulk expression of ANGPT1 in protoplasmic and fibrous astrocytes is displayed. **(c)** Dot plot showing ANGPT2 dysregulation in BEC and SMC sub-cell types, other cell types do not show significant dysregulation of this gene. Dot size represents the significance (negative decadic logarithm of the p-value), and the color the log fold change. **(d)** Dot plot of ANGPT1 dysregulation showing only significant upregulation for fibrous and protoplasmic astrocytes. Dot size and color follow the same logic as in (c). **(e)** Spearman-correlation matrix between ANGPT1 and ANGPT2 expression across cell types. Cell types such as BEC are separated by horizontal and vertical solid lines. The color scale represents the directionality and quantity of correlation. **(f)** Symbolic intersections of ANGPT1 and ANGPT2 dysregulation induced by overexpression of ANGPT2 BECs and SMCs and additional upregulation of TIE2/TEK in BECs, followed by overexpression of its antagonist ANGPT1 in astrocytes**. (g)** Spatial-transcriptomics slices from male persons labelled as control (left column) and severeAD (right column, as defined by Gong et al.^44^) highlighting the spots expressing either ANGPT1, ANGPT2, or both (>0 normalized expression of the corresponding gene is defined as expressed). The black line estimates the white and grey matter border defined by the MBP oligodendrocyte marker gene (bottom row), i.e. high expression white matter, low expression grey matter. **(h)** Density plot of ANGPT2 expression, in control and severeAD omitting the zero-entries.

To quantify these associations, we compared the highest versus lowest tertiles directly. This confirmed that *ANGPT2* expression was significantly higher in brain endothelial cells (BECs) and smooth muscle cells (SMCs) of individuals with high NP counts and high β-amyloid load (***Fig. 3c***). The association was particularly strong in all BEC subtypes from the high NP group (1.15 < logFC < 1.28, p < 10⁻⁸⁰) (***Supplemental Table 3***). Conversely, higher *ANGPT1* expression was observed specifically in protoplasmic and fibrous astrocytes of individuals with high NP and neurofibrillary tangle (NFT) burden (***Fig. 3d***).

To understand the relationship between these opposing cell-type patterns, we correlated pseudobulk expression levels across all 101 individuals (***Fig. 3e, Supplemental Fig. 8***). This revealed a negative correlation between *ANGPT1* and *ANGPT2* within vascular cell types, such as between *ANGPT2* in capillary BECs and *ANGPT1* in arteriolar SMCs. Interestingly, we also observed positive correlations, including between in protoplasmic astrocytes and arterial SMCs. These findings support a model where pathological *ANGPT2* expression from BECs and SMCs is countered by a potentially compensatory *ANGPT1* response from neighbouring astrocytes (***Fig. 3f***). To validate this in a spatial context, we analysed an independent spatial transcriptomics dataset^44^. This analysis confirmed the co-expression of *ANGPT1* and *ANGPT2* in the cortex and revealed a significant elevation of *ANGPT2* expression in pathological AD cases compared to controls (p < 2.22x10⁻¹⁶) (***Fig. 3g,h***) and could be shown in other datasets as well (***Supplemental Fig. 9 and 10***).

Given the prominent role of angiopoietins in vascular remodeling, our findings point to prominent dysregulation of angiogenesis and support a model whereby anti-angiogenic signals (*ANGPT2*) from BECs associated with β-amyloid and NPs compete with pro-angiogenic signals (*ANGPT1*) from astrocytes.

### Lesion specific dysregulation BBB-linked pathways and cell proliferation

To refine the relationship between cell type specific gene expression and pathological lesions, we computed associations between gene expression and biological pathways and illustrated them in DicePlots^45^ **(*Fig. 4a, Supplemental Figure 11, Supplemental Table 4)***. This visualization pointed to three broad groups of pathways: First, a group of BBB-linked pathways, such as focal adhesion and cell-substrate junction, were significantly dysregulated across all neuropathological lesions in multiple cell types and most prominently in venous BEC, protoplasmic astrocytes, and M-pericyte. Second, we observed a specific association between high NP counts and dysregulated cell proliferation pathways in vascular cells but not in microglia or astrocytes. Third, glial cells showed distinct pathway enrichments, with microglia and protoplasmic astrocytes being particularly responsive to high neurofibrillary tangle (NFT) density.

**Fig. 4.**
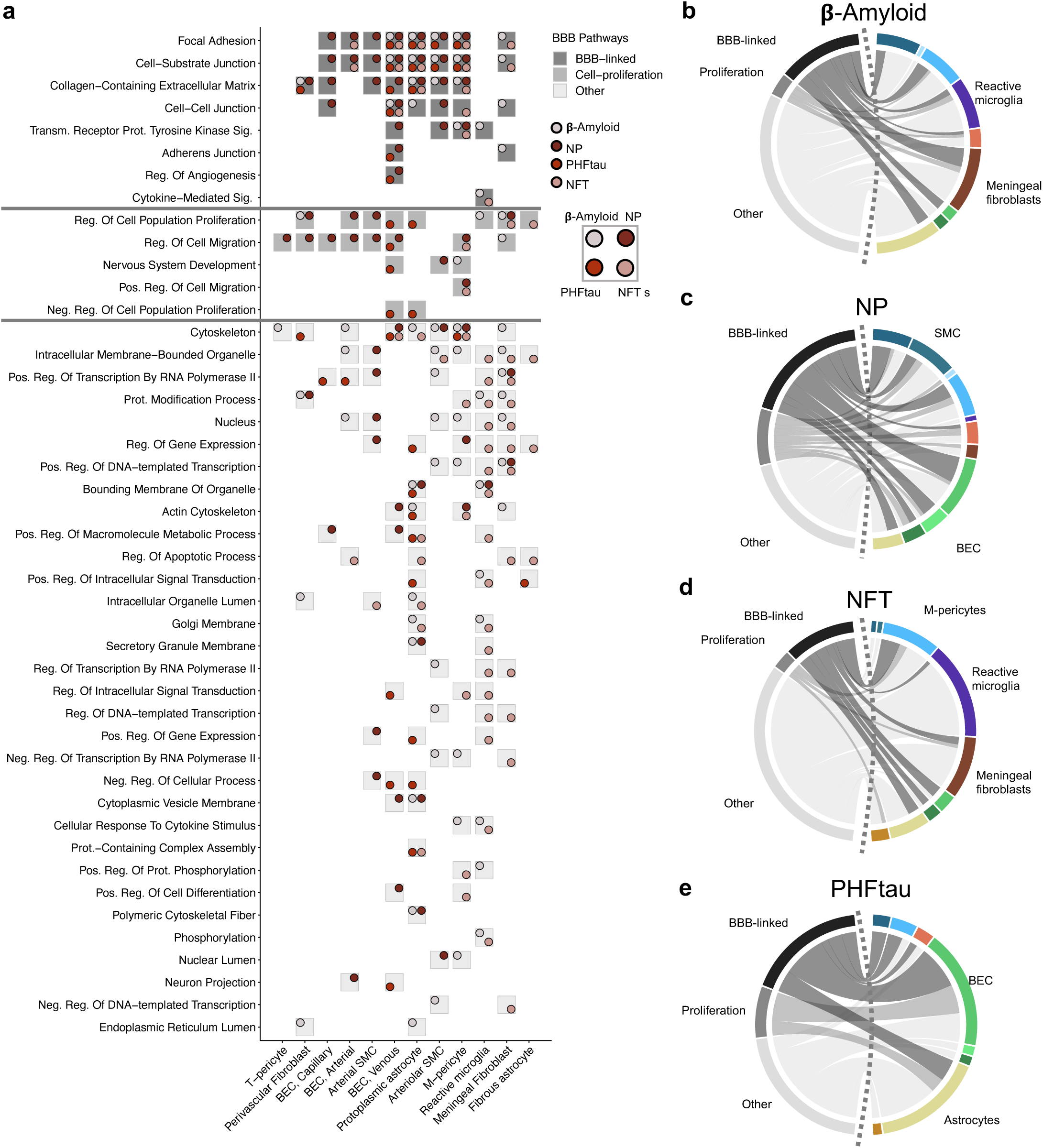
Pathway analysis across neuropathological lesions and plaques in vascular niche: **(a)** DicePlot for gene ontology pathways (KEGG and Reactome pathways are presented in the supplemental material) and cell types for the different neuropathological burdens. Each cell contains between 1 and four dots. These dots represent significant association of the pathway in the cell type for the four neuropathological burdens defined on the right side. BBB-linked, proliferation and other pathways are highlighted by different shades of gray and separated by horizontal lines. **(b-e)** Chord diagram displaying the proportions of dysregulated BBB-linked and non-BBB-linked pathways to the different cell types in β-amyloid **(b)**, NP **(c)**, NFTs **(d)**, and PHFtau **(e)**.

Aggregating these dysregulated pathways provided a global view of which cell types were most affected by each specific lesion **(*Fig. 4b-e*)**. The likely pathological progression from β-amyloid to NPs, tangles, neurofibrillary tangles reveal distinct cellular vulnerabilities. High β-amyloid load was predominantly associated with pathway dysregulation in reactive microglia, meningeal fibroblasts, and protoplasmic astrocytes (***Fig. 4b***). In contrast, high NP counts were linked most strongly to pathways in BECs and SMCs (***Fig. 4c***). High NFT density was associated with responses in microglia, meningeal fibroblasts, and M-pericytes (***Fig. 4d***), while high PHF-tau load was linked to dysregulation of BECs and protoplasmic astrocytes (***Fig. 4e***). Overall, these analyses reveal that amyloid and tau pathologies are associated with surprisingly distinct patterns of pathway dysregulation across specific vascular and glial cell types.

### Increasing molecular correlation between cognitive decline and neuropathological burden in progressing AD

A known discrepancy exists between the burden of AD neuropathology and the severity of clinical symptoms. To investigate the molecular basis of this, we compared transcriptional signatures associated with neuropathological scores to those associated with clinical diagnoses of no cognitive impairment (NCI), mild cognitive impairment (MCI), and AD dementia (***Fig 5a***). Pairwise comparisons revealed that the largest transcriptional differences were between the MCI and AD groups versus the NCI group, with far fewer DEGs found in the direct AD versus MCI comparison (***Fig. 5b, Supplemental Table 5***). Specifically, the comparisons with the NCI group showed far more significant DEGs than that between MCI/AD (MCI/NCI: 1,558; AD/NCI: 2,084; AD/MCI: 586). This pattern was observed across all cell types, except in microglia, where similar numbers of DEGs were found between NCI/MCI (34 DEGs) and MCI/AD (36 DEGs) comparisons. Overall, these data indicate that transcriptional signatures of vascular dysfunction emerge early at emerge at the mild cognitive impairment (MCI) stage, preceding severe cognitive symptoms and aligning more closely with AD than cognitively normal individuals.

**Fig. 5.**
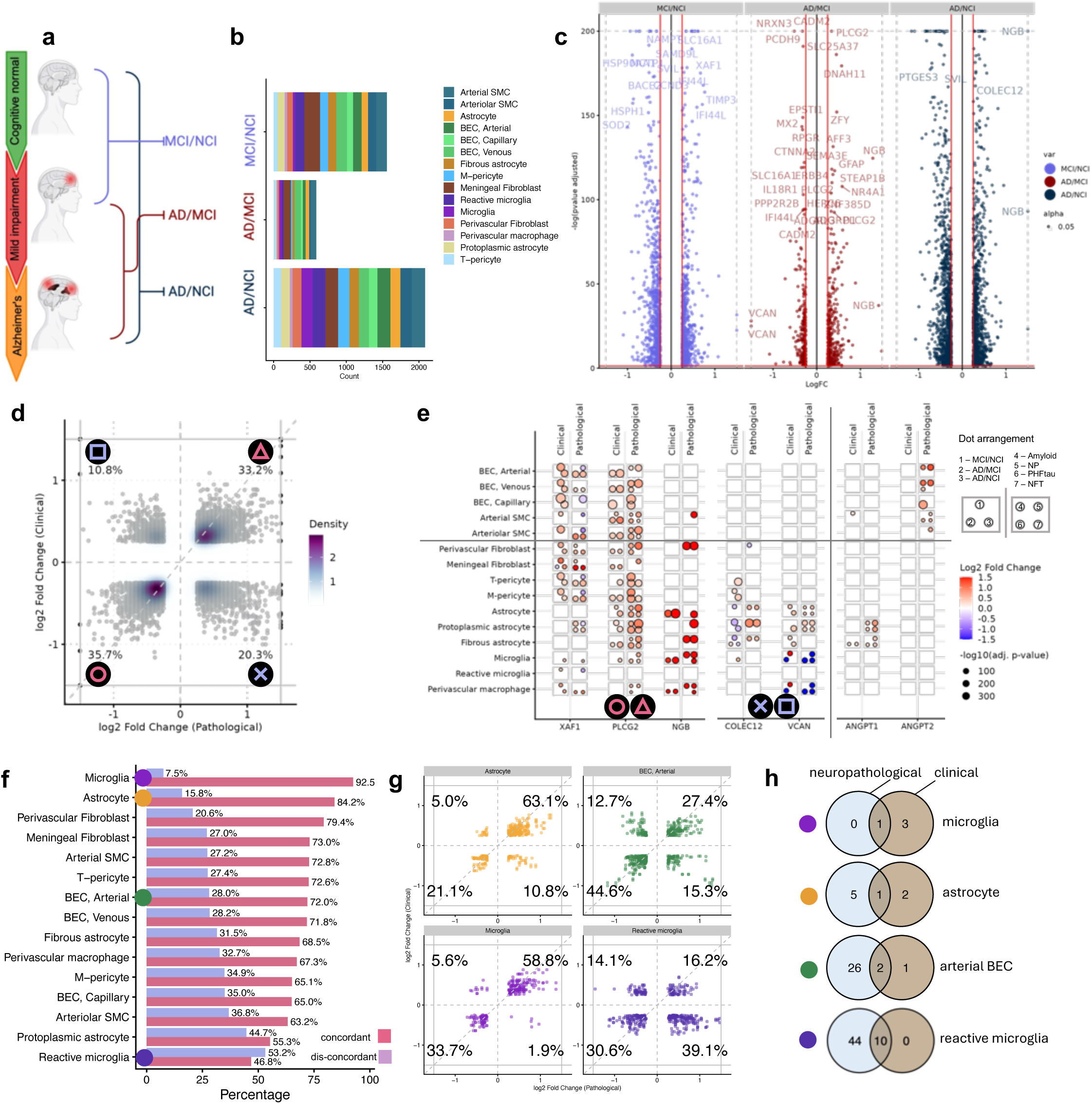
Alignment of molecular patterns related to cognitive decline and to neuropathological burden: **(a)** Schematic view of the clinical diagnosis for no cognitive impairment (NCI), mild cognitive impairment (MCI), and Alzheimer’s disease (AD) and the derived comparisons for computing DEGs. **(b)** Bar plot displaying the number of DEGs by clinical diagnoses comparison, colored by fine cell type. **(c)** Volcano plots presenting the log fold change and negative decade logarithm of the genes in the three comparisons. Selected genes are highlighted. **(d)** The plot presents genes that are up- or down-regulated in both the clinical and neuropathological definition. Genes that are differentially regulated in only one of the considerations are not shown (white horizontal / vertical area). Each gene can be present multiple times because it is included in several cell types. In addition to the single points, a density representation is shown in the background. The percentages of genes irrespective of the cell type are presented in the four sectors and represented by four different symbols. **(e)** DicePlot presenting for selected genes from (d) in the high-resolution view. For each gene the dysregulation in the 15 cell types is presented. For each combination of a gene and cell type, each box contains up to seven dots representing three clinical comparisons (AD/NCI, MCI/NCI, AD/MCI) and four neuropathological burdens (high versus low β-amyloid, NP, tangles and NFT). If no dot is presented, the gene is not dysregulated for the respective comparison in the respective cell type. Symbol(s) below the gene describe from which area of the scatter plot in panel (d) the gene was derived. Note that a gene can be included in several plot areas because it might be concordant in one but discordant in another cell type. **(f)** Percentage of concordant versus discordant genes for each of the 15 cell types. Four cell types with high, medium and low concordance are presented as scatter plot in panel (g). **(g)** Like panel (d), the genes are now presented for four different cell types. Coloring matches the cell type defined color consistently used in the manuscript. Percentages in the corners denote the fraction of genes per area. **(h)** Venn diagrams presenting the overlap of significant pathways between the clinical and neuropathological definitions for the cell types from (f) and (g). In microglia, pathways are dominantly significant in the clinical definition for the other cell types relying on the neuropathology.

To understand the functional differences between gene signatures tied to pathology and cognition, we next compared the trajectories of key DEGs across these two contexts. While genes previously implicated in AD like *XAF1* and *PLCG2* were among the DEGs identified in clinical comparisons, the analysis revealed a striking divergence for the top pathology-associated genes (***Fig. 5c, Supplemental Table 5***). Specifically, *VCAN* and *NGB*—two genes with the largest fold-changes in the pathology analysis—showed differential expression only with late-stage cognitive decline. Similarly, *ANGPT1* and *ANGPT2* were not prominently correlated with clinical disease. These results demonstrate a clear divergence, indicating that the most robust molecular markers of neuropathology are not the same as those that mark clinical disease status.

To systematically quantify this divergence at cell type resolution, we directly compared the direction of gene expression changes across both analyses. While nearly 70% of DEGs were concordant overall (***Fig. 5d***), a detailed view of select genes like *COLEC12* and *VCAN* highlighted discordant patterns in specific cell populations (***Fig. 5e, Supplemental Fig. 12***). A global assessment of concordance for each cell type confirmed that the degree of alignment is highly cell-type specific (***Fig. 5f***). Microglia showed the highest concordance (92.5%), followed by astrocytes (84.2%) and perivascular fibroblasts (79.4%) (***Fig. 5f,g, Supplemental Fig. 12, 13***).

To understand the broader biological implications of these findings, we compared enriched pathways between the clinical and pathological contexts. Our analysis revealed fewer significant pathways associated with clinical status (67 total hits, representing 24 different pathways) compared to neuropathology (574 total hits representing 127 different pathways) (***Supplemental Figure 14***). However, pathway significance varied by cell type (***Fig. 5h; Supplemental Fig. 11***). In astrocytes, arterial BECs, and reactive microglia we identified more pathways based on neuropathological burdens than those based on cognition. In contrast, no pathway was significant in microglia based solely on pathology; instead, three pathways, including nervous system development and regulation of gene expression, were significantly associated with cognition, particularly in the AD versus NCI comparison. This identification of cognitively-associated microglial pathways underscores the role of the immune system in cognitive decline along AD progression.

## DISCUSSION

This study addresses the critical role of the neurovascular unit in Alzheimer’s disease (AD) by generating a deep, cell-type-resolved transcriptomic atlas from 101 individuals using our VINE-seq methodology. Our analysis reveals that vascular dysfunction in AD is driven primarily by profound gene dysregulation within resident cells rather than by changes in their relative proportions, with brain endothelial cells (BECs) and smooth muscle cells (SMCs) exhibiting the most pronounced transcriptional changes. These molecular signatures emerge early in the clinical course; the transcriptional profile of MCI already aligns more closely with AD than with cognitively normal individuals. Further, we demonstrate that amyloid and tau pathologies are associated with distinct cellular responses: amyloid is predominantly linked to dysfunction in vascular cells, a process hallmarked by the dysregulation of angiopoietin signalling, while tau pathology is more strongly linked to responses in glial cells. Together, these findings decompose the general concept of “vascular dysfunction” into specific, cell-type- and lesion-dependent molecular programs that arise early in AD.

A key molecular finding of our study is the identification of a pathological push-pull dynamic within the angiopoietin signalling axis, driven by opposing expression patterns of *ANGPT1* and *ANGPT2*. While elevated circulating angiopoietins have been linked to AD and blood-brain barrier (BBB) leakiness in prior studies^46–48^, our work provides the first high-resolution cellular map of this process in the human brain. We pinpoint vascular cells (BECs and SMCs) as the source of pathological *ANGPT2* associated with amyloid burden, which is countered by a potentially compensatory upregulation of the vascular-stabilizing factor *ANGPT1* from neighbouring astrocytes. This dynamic, validated in independent single-cell and spatial datasets, suggests that a central feature of AD neuropathology is a competition between amyloid-induced vascular destabilization and an astrocyte-led attempt to preserve vascular integrity.

Our comparison of vascular transcriptional patterns associated with pathology versus those linked to cognitive decline revealed a profound divergence between the two. We demonstrated that the most robust molecular signatures of AD pathology, including the dysregulation of BBB-linked pathways, are not direct correlates of cognitive impairment. In contrast, microglia exhibited strong concordance in their transcriptional state between pathology and cognition. This suggests that while vascular dysfunction is a direct molecular consequence of accumulating lesions, it is the brain’s innate immune response, orchestrated by microglia as they react to this altered and likely compromised brain environment, that more closely determines the clinical trajectory. This positions cells of the neurovascular unit as key regulators of microglial homeostasis^49,50^, whose dysregulation then governs the progression from pathological burden to cognitive impairment.

In conclusion, by generating a cell type-resolved atlas of the neurovascular unit along AD progression, this study provides new insights into the key molecular pathways mediating vascular dysfunction. We demonstrate that the molecular changes associated with AD are not uniform but are highly specific to both cell type and pathological lesion, and that these changes are established early at the MCI stage. Crucially, our work disentangles the early molecular signatures of vascular dysfunction, such as dysregulation of the angiopoietin axis, from the downstream immune responses that more closely align with cognitive decline. This foundational resource paves the way for future research to untangle these distinct mechanisms and may inform the development of novel diagnostics and therapies targeting either the initial vascular insults or the subsequent glial responses that drive the disease.

### Limitations

While our cohort of 101 participants provides a deep view into the neurovascular unit, we acknowledge several limitations. First, as with any single-cohort study, our findings warrant replication in independent and diverse populations to ensure generalizability. Second, the use of our VINE-seq protocol, while essential for enriching vascular nuclei, may introduce cell-type biases inherent to enrichment-based approaches and complicates direct comparisons with studies using different methodologies. Finally, we acknowledge that the different neuropathological measures (e.g., β-amyloid load, NP counts) are themselves correlated in AD cases, meaning the sets of differentially expressed genes (DEGs) derived from them are not statistically independent. Despite these considerations, this dataset represents a valuable and comprehensive resource for the research community, enabling fellow researchers to validate findings and generate new hypotheses about the role of the vasculature in AD.

## METHODS

### Case selection from ROSMAP

We selected 101 individuals from the Religious Orders Study or Rush Memory and Aging Project (ROSMAP), ongoing prospective cohort studies of brain aging and dementia^34^. ROSMAP participants enroll without known dementia and agree to annual clinical evaluation and brain donation. Informed and repository consents and an Anatomic Gift Act was obtained from each subject. Both studies were approved by an Institutional Review Board (IRB) of Rush University Medical Center. Group characteristics and metadata are presented in **Supplemental Table 1**. Individuals are classified as Alzheimer’s dementia, mild cognitive impairment (MCI), or no cognitive impairment (NCI)^12^ NIA-Reagan was used for a pathologic diagnosis of AD^10^. We also examined neuritic plaques (NP). AD was also quantified with molecularly specific markers for β-amyloid load and PHFtau tangles. Methods are previously reported^10,12,40–42,51^. Brain multi-omic data is available from the same persons as previously reported^16^.

### Isolation and single-cell sequencing of vascular nuclei from post-mortem brain tissue

All procedures were carried out as previously reported^52^, on ice in a 4 °C cold room as rapidly as possible. Briefly, brain tissue (0.4 grams or more) was thawed on ice for 5 min with 5 ml of nuclei buffer (NB): 1% BSA containing 0.2 U μl−1 RNase inhibitor (Takara, 2313A) and EDTA-free protease inhibitor cocktail (Roche, 11873580001). Tissue was quickly minced and homogenized with 7-ml glass douncers (357424, Wheaton). Homogenates were vigorously mixed with chilled dextran (D8821, Sigma) to a final volume of 18% dextran before centrifugation at 4,400g for 20 min with no brake. After centrifugation, samples separate into a top myelin layer, middle parenchymal layer and vascular-enriched pellet. The myelin layer was aspirated, tips were changed, and the parenchymal layer was carefully removed without disturbing the pellet. Vascular-enriched pellets were gently resuspended in 1 ml of NB and added to pre-wetted 40-μm strainers sitting on top of 50-ml falcon tubes. Strainers were washed with 100 ml of cold PBS before mashing vascular fragments through the cell strainer using the plunger end of a 5-ml syringe, with elution via 40 ml of PBS. Liberated vascular cells were pelleted, filtered through a 40-μm strainer (Flowmi), transferred to FACS tubes, stained with Hoechst 3342 (1:2,000, Thermo Fisher Scientific) and rabbit monoclonal anti-NeuN Alexa Fluor 647 (1:500, Abcam, ab190565), and nuclei collected on a SH800S Cell Sorter into chilled tubes containing 1 ml of NB without protease inhibitor. Libraries were prepared using the Chromium Single Cell 3ʹ v3.1 according to the manufacturer’s protocol (10x Genomics). Fifteen PCR cycles were applied to generate cDNA before 16 cycles for final library generation. Generated snRNA-seq libraries were sequenced on S4 lanes of a NovaSeq 6000 (150 cycles, Novogene).

### Bioinformatics and data analysis

#### Single-nucleus RNA-seq data processing and quality control

Raw sequencing samples were processed using the CellRanger^53^ software v7.0.0 by 10X Genomics, counting both intronic and exonic reads during mapping against the provided reference genome GRCh38-2020-A. Technical replicates from multiple sequencing batches were pooled for each sample. The number of expected cells / nuclei per sample was set to 5,000. Resulting raw barcode to gene count matrices were then analyzed with SoupX^54^ v1.6.0 at standard parameters to correct for contaminating ambient RNA and perform cell / nuclei calling. Filtered barcode to gene matrices were then loaded into Seurat^55^ v4.1.1 objects, removing low-quality genes and cells by requiring at least 5 detected cells and 200 features, respectively.

#### Doublet detection and removal

Following the standard Seurat workflow with scTransform v0.3.5 and 20 principal components set, we applied DoubletFinder v3 to each sample to remove contaminating doublet and multiplet nuclei. Thereby, the core parameters of DoubletFinder were selected as follows: pN=0.25, doublet rate = 0.15, PCs=20. We followed the officially recommended way to auto-estimate the remaining parameters pK and nExp for each sample, based on a parameter sweep, i.e. grid search. Using SeuratDisk v.0.0.0.9020 all Seurat objects were computed from file format .h5seurat into .h5ad on-disk. We then created a combined anndata object with Scanpy^56^ v1.9.1 by concatenating all count matrices with an outer join on the features, excluding all cells that were flagged as “doublet” by DoubletFinder^57^. From the combined matrix, all cells and features, with less than 250 detected features and 50 detected cells, respectively were further discarded. Furthermore, nuclei with less than 7000 expressed genes in total and a mitochondrial content of less than 5% of the entire transcriptome were retained only.

#### Data integration and dimensionality reduction

Following the standard analysis workflow of Scanpy we determined the top 3,000 highly variable genes, normalized as well as scaled the counts and performed dimension reduction with PCA (Principal Component Analysis) and UMAP using 11 significant principal components. Data integration was performed on a Nvidia A100 using the Pytorch-based GPU implementation of Harmony^58^ through Harmonypy v0.0.6 and Harmony-pytorch v0.1.7, with the donor sample set as batch key across the nuclei and a maximum of 25 iterations. The subsequently required distance graph was computed for n=100 neighbors using a GPU-implementation provided by the rapidsai^59^ package v22.06.00 on the harmonized data representation. Similarly, the global Uniform Manifold Approximation and Projection (UMAP) representation was determined using a minimal distance of 0.5, a spread of 0.8 and with an initial spectral clustering via rapidsai.

#### Clustering and cell type annotation

Cell clusters were called using a GPU-implementation of the louvain algorithm through rapidsai, initially setting the resolution to 0.55. Iteratively, we determined cluster markers, annotated cell populations using previously reported markers as reference. One low-quality cluster showing ambiguous marker expression and thus of undetermined origin was subsequently removed. Finally, we excluded nine samples having too few high-quality cells left for analysis (individual IDs C14, C15, C16, A11, C19, A20, M2, M14, and M16). We aimed to further remove potentially undetected heterologous doublets from our combined dataset by defining a cell type-specific list of contaminating marker genes originating from the other cell types (see Supplemental Table 7). We then discarded all cells having 4 or more raw counts across these gene panels using their cell type specific list. We repeated the integration procedure to determine the final neighbor graph and nuclei embedding dimensions, reannotating the major cell type clusters. To generate a fine-grained (sub) cell type annotation, we subset nuclei from each cell type cluster and reperform the variable gene identification, count normalization, scaling, harmonizing, neighbor graph calculation, dimension reduction, Louvain cluster calling and cluster marker identification workflows. Again, using previously reported sets of markers we manually performed a detailed annotation of the resulting clusters. The procedure yielded our final data object with 842,646 high-quality nuclei and RNA counts across 29,871 human gene transcripts. Using the SeuratDisk package, we then converted the h5ad file instance back into a h5Seurat object, required for downstream analysis. We then clustered the gene-wise Spearman correlation matrix derived from normalized z-scores for the globally top 5,000 highly variable genes. We identified a cluster of N=754 highly correlated genes across the complete set of nuclei, containing also highly expressed neuronal markers, which we excluded from the feature matrix for all our downstream analysis. To study differential expression, we determined DEGs both on the single-nucleus- and sample-level by using MAST^60^ through Seurat. We used the covariate person sex as a latent variable to be regressed out during model generation. For assessing the dysregulation of genes concerning the neuropathological response variables we computed the DEGs based on the low and high tertile. As in the previous analysis we used MAST with “sex” as a latent variable for computation. Besides the 754 highly correlated genes, we exclude all non-coding RNAs from downstream analysis in a final filtering step.

#### Pathway Analysis

We use EnrichR^61^ for computing the enriched pathways and selected “KEGG_2021_Human”, “Reactome_2022”, “GO_Biological_Process_2023”, and “GO_Cellular_Component_2023” for computing the pathways. For computation, only DEGs with an adjusted p-value smaller 0.05 have been considered with absolute logFC>0.25. The pathways categories were manually curated into the groups “BBB-linked”, “Proliferation-linked”, and “Other” (***Supplemental Table 6***). We omit pathways that occur only once from displaying in the DicePlots (***Supplemental Fig. 11***).

#### Validation of findings using Mathys 2023 and cellxgene census

Mathys et al.^16^ DEG results were downloaded from https://github.com/mathyslab7/ROSMAP_snRNAseq_PFC using the global pathology DEG contrast. Moreover, we use the CELLxGENE cell-census (census_version=”2023-12-15”) to confirm our DEG findings. For this we used the following filtering: We only considered cells from individuals older or equal to 40 years, and in the following conditions: normal, dementia, Parkinson’s disease, and Alzheimer’s disease. The cells underwent QC, where all cells with >= 5% mitochondrial counts, n_total_counts > 50K, or n_genes_by_counts > 6000 were excluded. Finally, we used the scVI framework to integrate the data with the batch key defined as dataset_id+assay+donor_id and by dynamically estimated epochs.

#### Spatial transcriptomics data analysis

We first converted the obtained GEF files (output of the Stereo-Seq Analysis Workflow (SAW) pipeline^62^) into Seurat objects for bin200 (200x200 Stereo-seq spots). This is done by using the *io.stereo_to_anndata* function in the Stereopy package^63^(v1.1.0), followed by applying the *h5ad2rds.R* script provided by BGI. After conversion, samples were normalized using the *NormalizeData* function in Seurat^55^ (v5.1.0) and subsequently merged using the *merge* function.

## Data availability

All data are available on https://synapse.org

All spatial transcriptomics data can be found on gene expression omnibus https://www.ncbi.nlm.nih.gov/geo/ GSE269906

All data used from CELLxGENE can be fetched directly using the provided code. The data used from Hahn et al. can be accessed using https://twc-stanford.shinyapps.io/spatiotemporal_brain_map/

ROSMAP resources can be requested at https://www.radc.rush.edu and/or https://www.synapse.org.

## Code availability

All code for primary analysis will be available upon publication.

Additionally, all code used for CELLxGENE analysis will be available upon publication.

## Acknowledgments

ROSMAP is supported by P30AG10161, P30AG72975, R01AG15819, R01AG17917, U01AG46152, and U01AG61356. K.A.Z. This work was supported by the Alzheimer’s Association Research Fellowship AARF-20-685030.

## Author contributions

**MF:** performed data analysis, wrote the manuscript draft**; AY:** performed experiments, wrote the manuscript draft**; FK:** contributed to data analysis**; SG:** performed spatial transcriptomics data analysis**; IFD:** contributed to data analysis**: HS: ; KAZ: ; DB: ; MRA: ; DC: ; SS: ; MAB: ; ECS: ; KB: ; FG:** contributed to data analysis**; DB:** contributed to the study design**; MB: ; AK:** supervised the data analysis, wrote the manuscript draft**; TWC:** supervised the study, wrote the manuscript draft. All authors made critical revisions to the manuscript.

## Materials & Correspondence

Correspondence and requests for materials should be addressed to Prof. Andreas Keller or Prof. Tony Wyss-Coray.

---

**Supplemental Figure 1:**
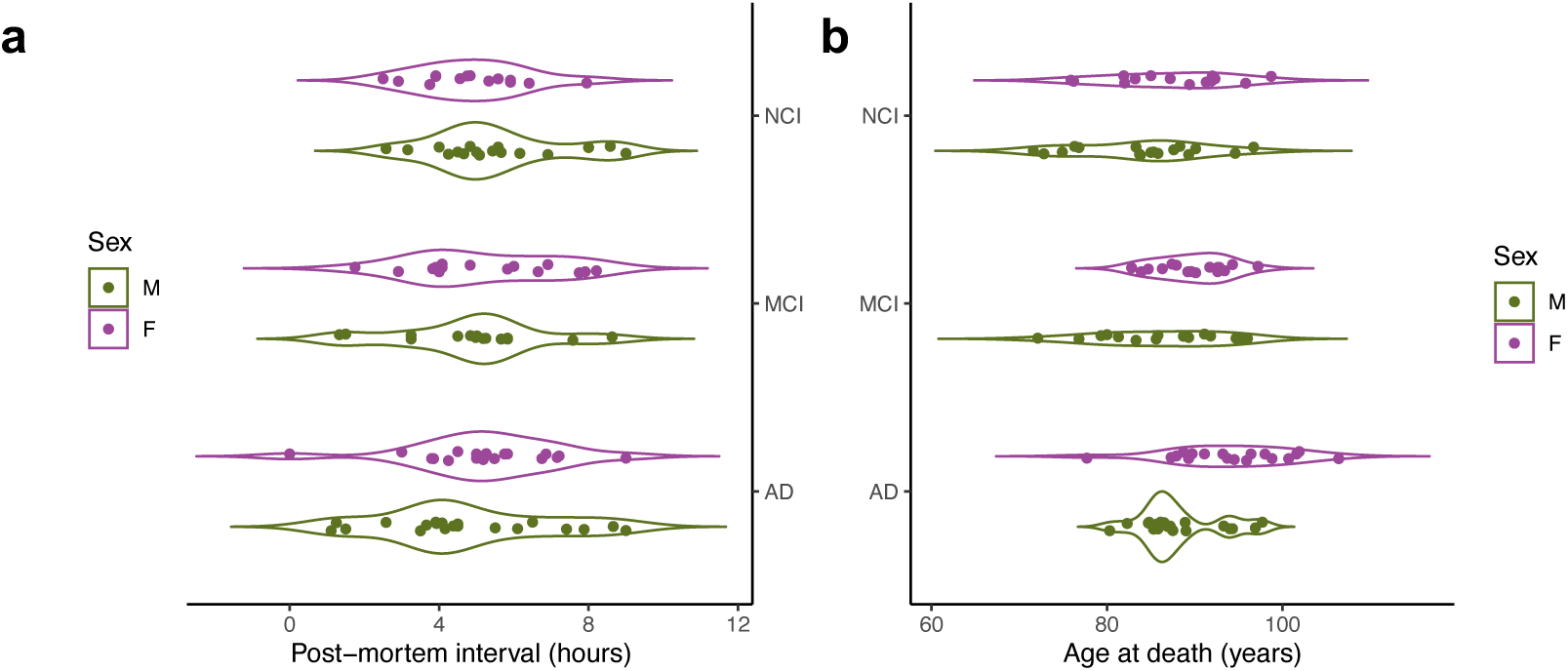
Demographic details (**a**) post-mortem interval, (**b**) age at death.

**Supplemental Figure 2:**
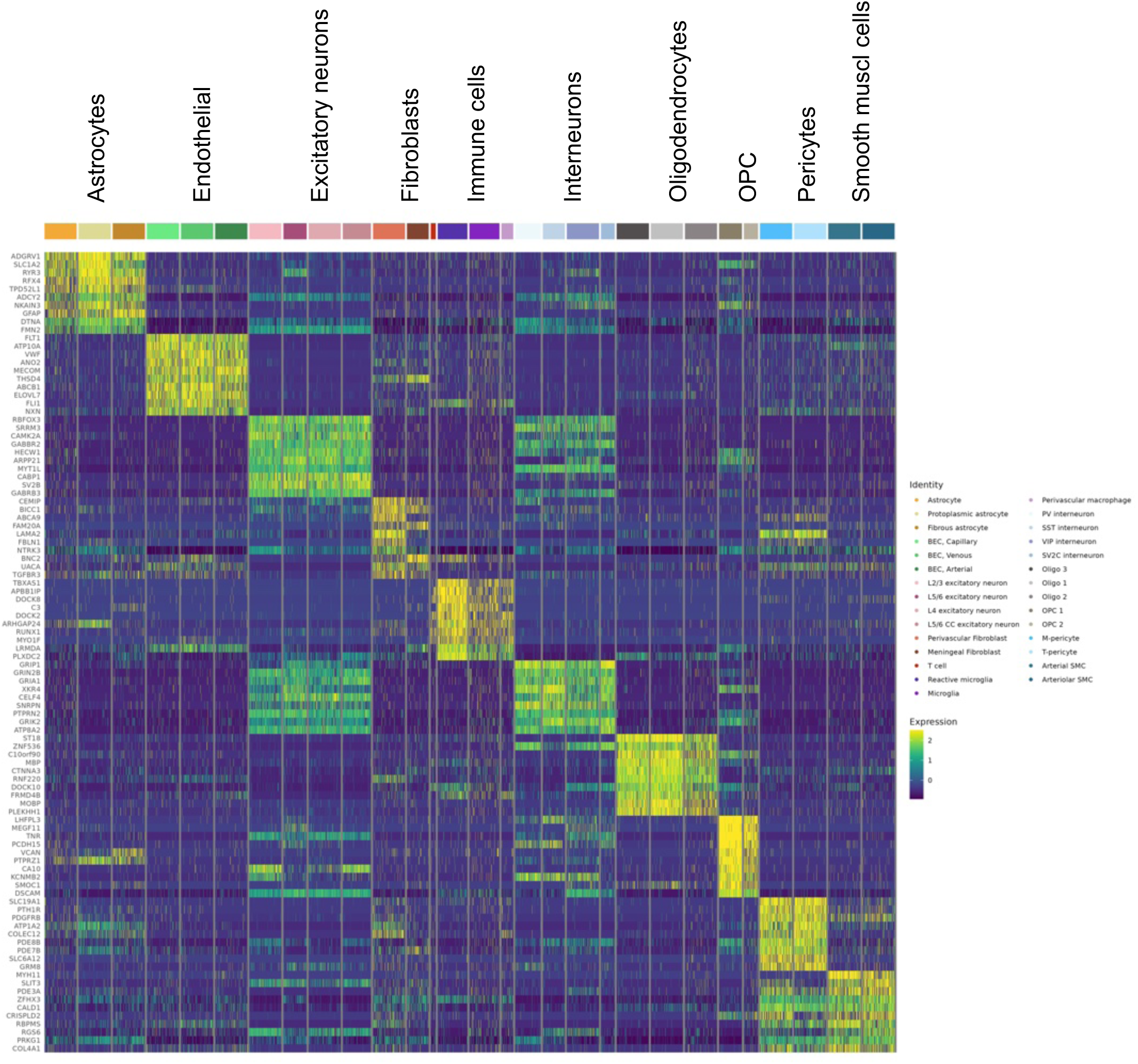
Broad cell type annotation heatmap, highlighting the top cell type marker.

**Supplemental Figure 3:**
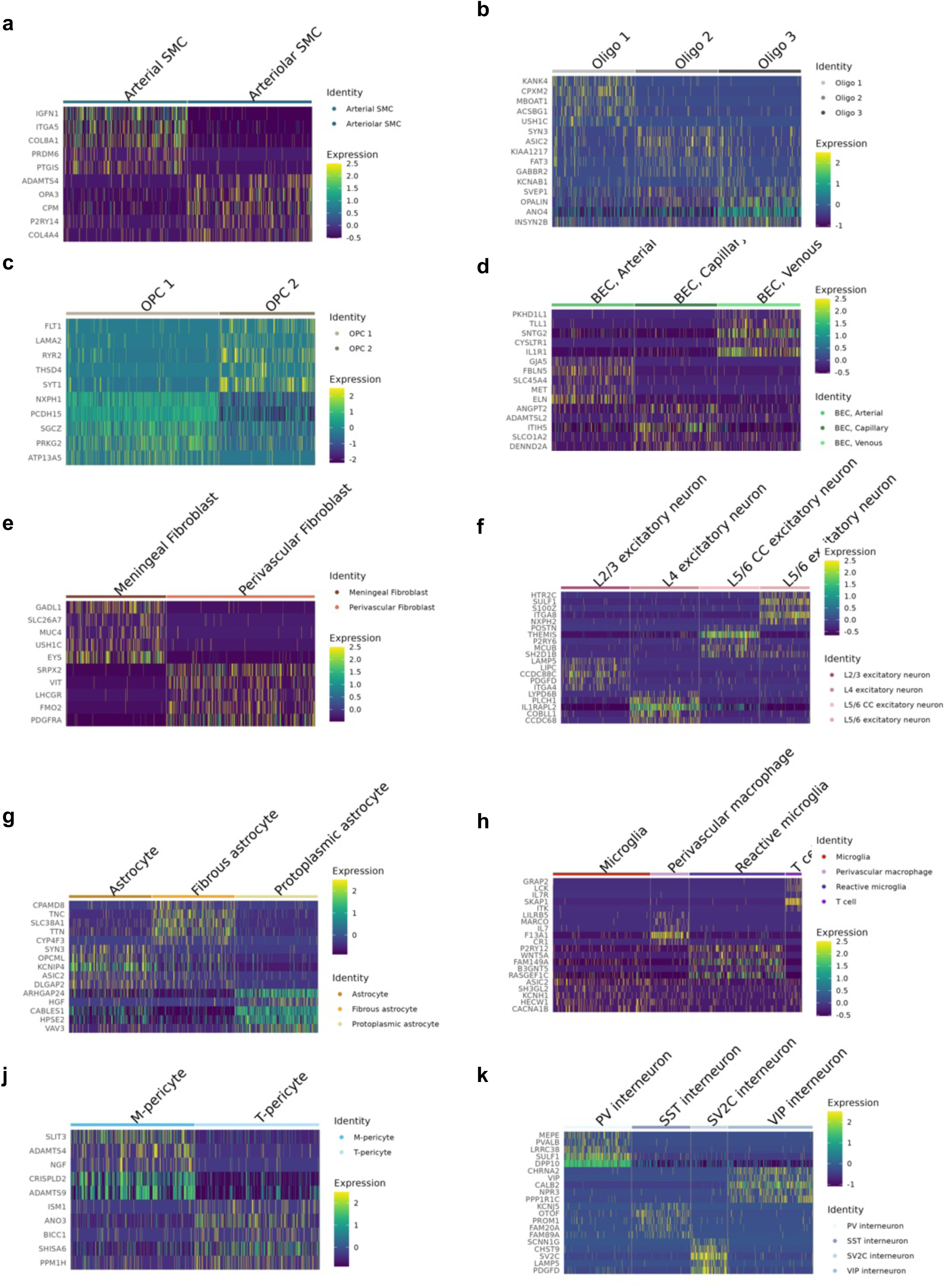
Fine cell type annotation highlighting the top marker for the fine cell type annotation for each broad cell type(**a-k**).

**Supplemental Figure 4:**
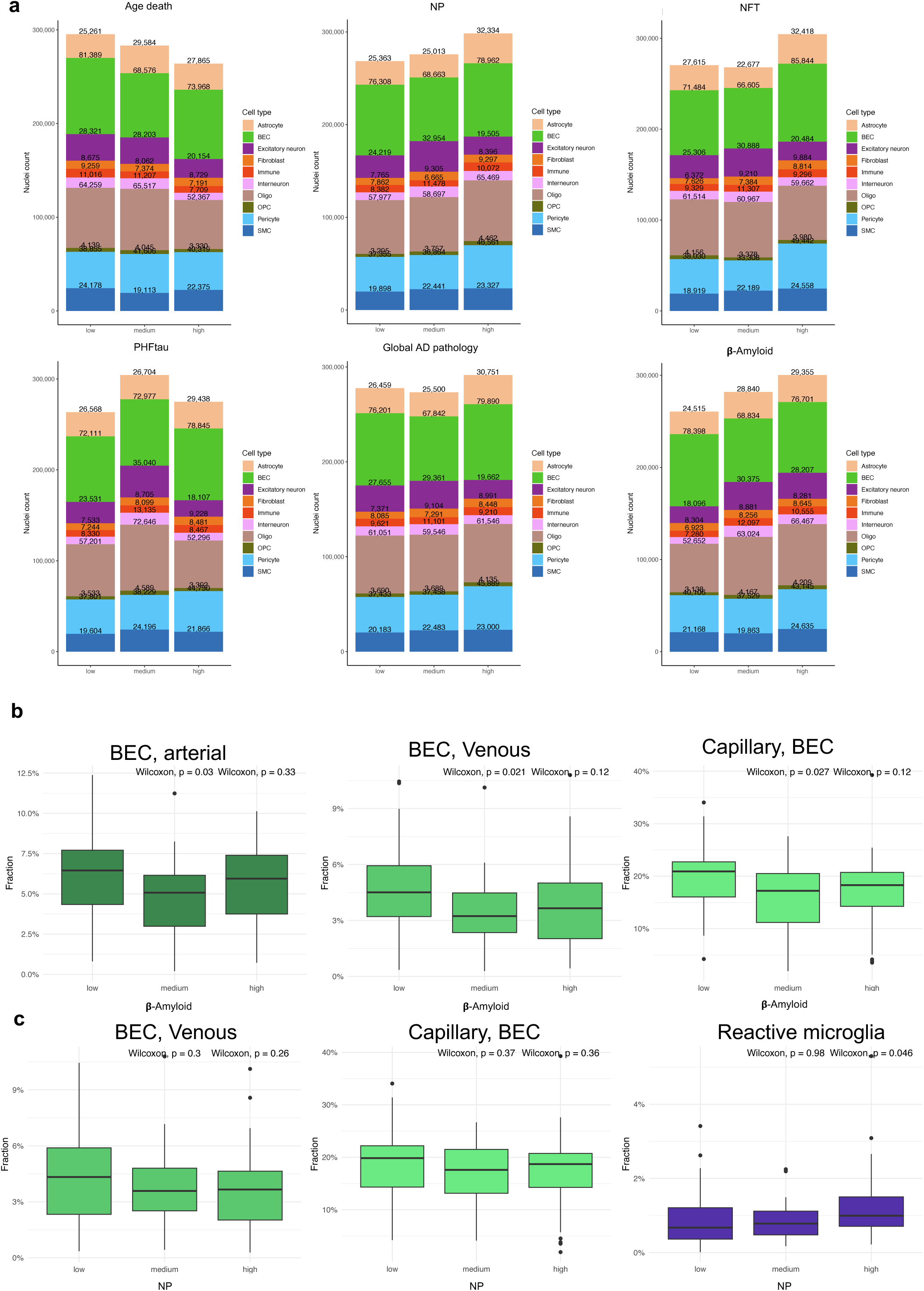
Compositional changes across metadata (**a**), and for BEC subtypes in β-amyloid tertiles.

**Supplemental Figure 5:**
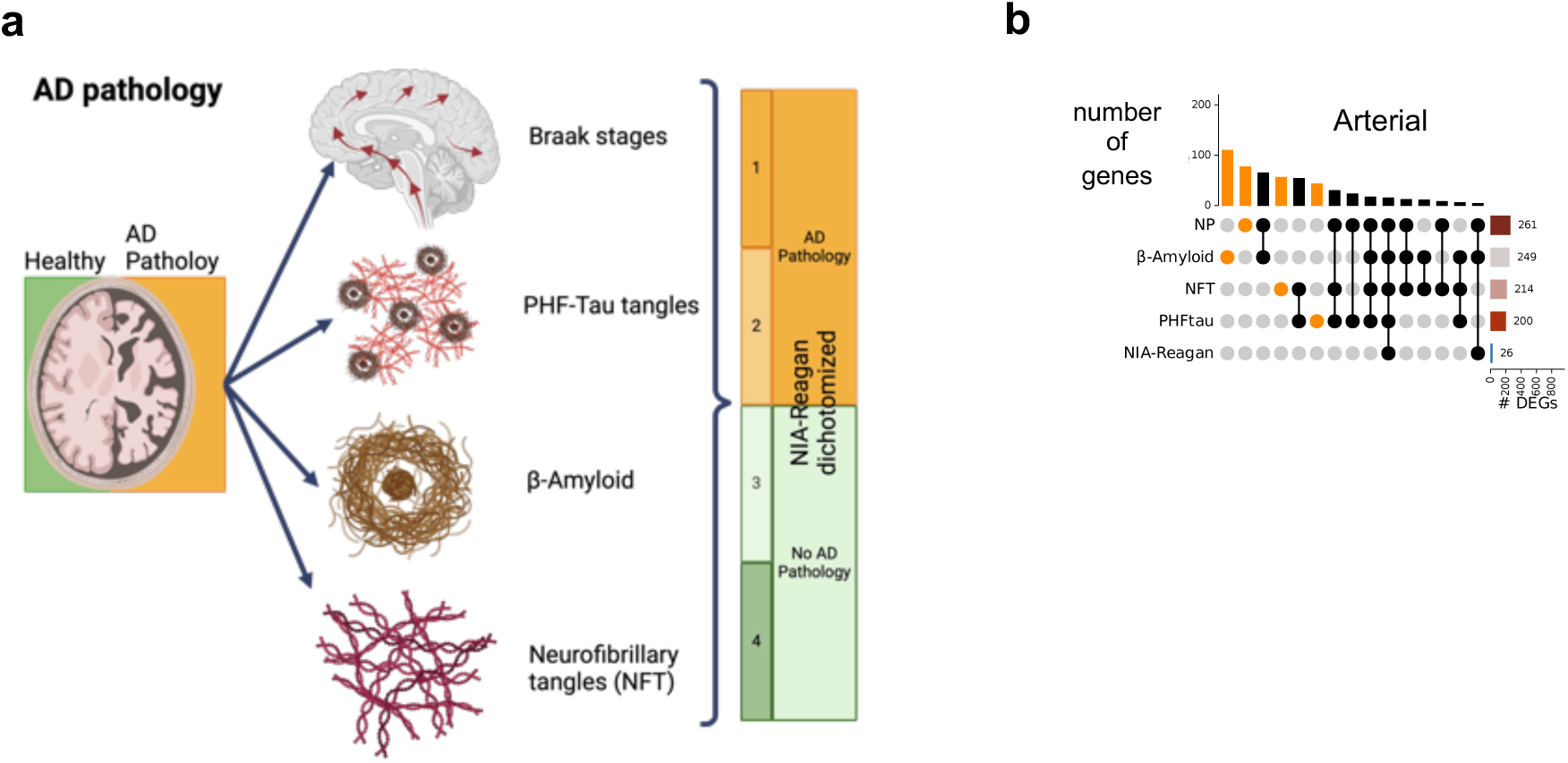
**(a)** Schematic of neuropathological lesions summarized into the NIA-Reagan score. Our analyses rely on the binarized NIA-Reagan score.

**Supplemental Figure 6:**
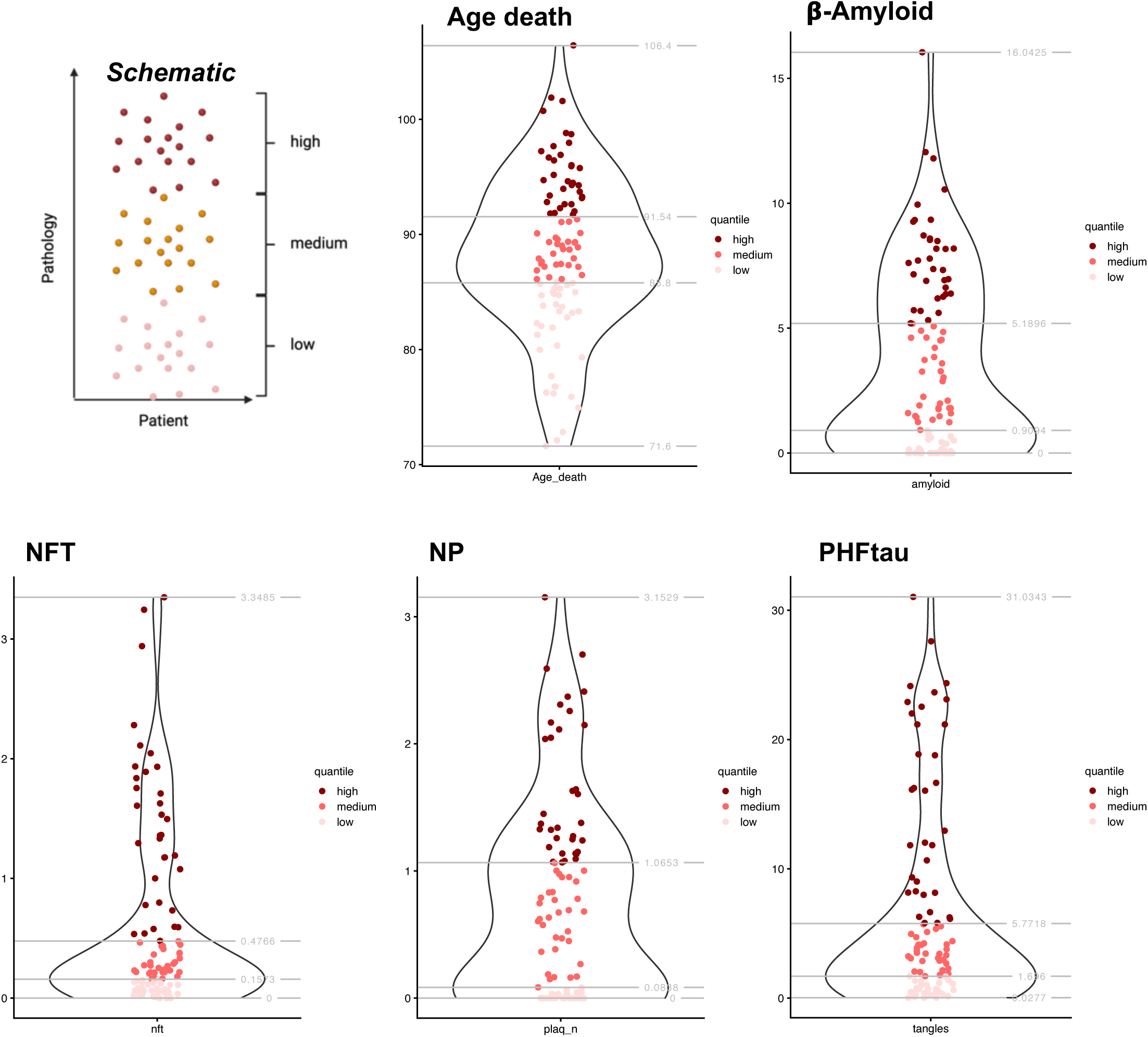
**(a)** Tertiles of the different plaques and meta information.

**Supplemental Figure 7:**
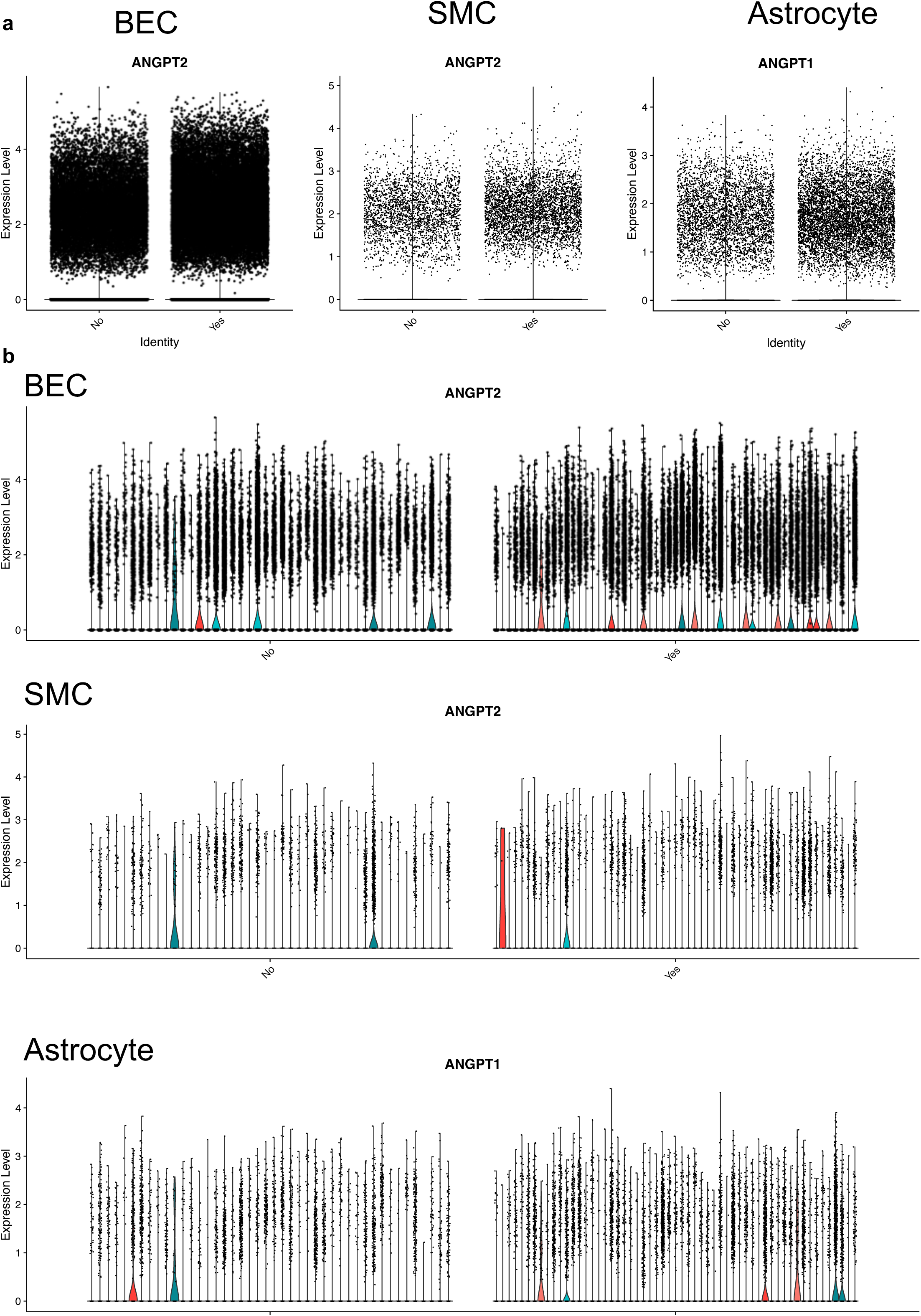
ANGPT1/2 expression by NIA-Reagan diagnosis, across all donors (**a**) and split by donors (**b**)

**Supplemental Figure 8:**
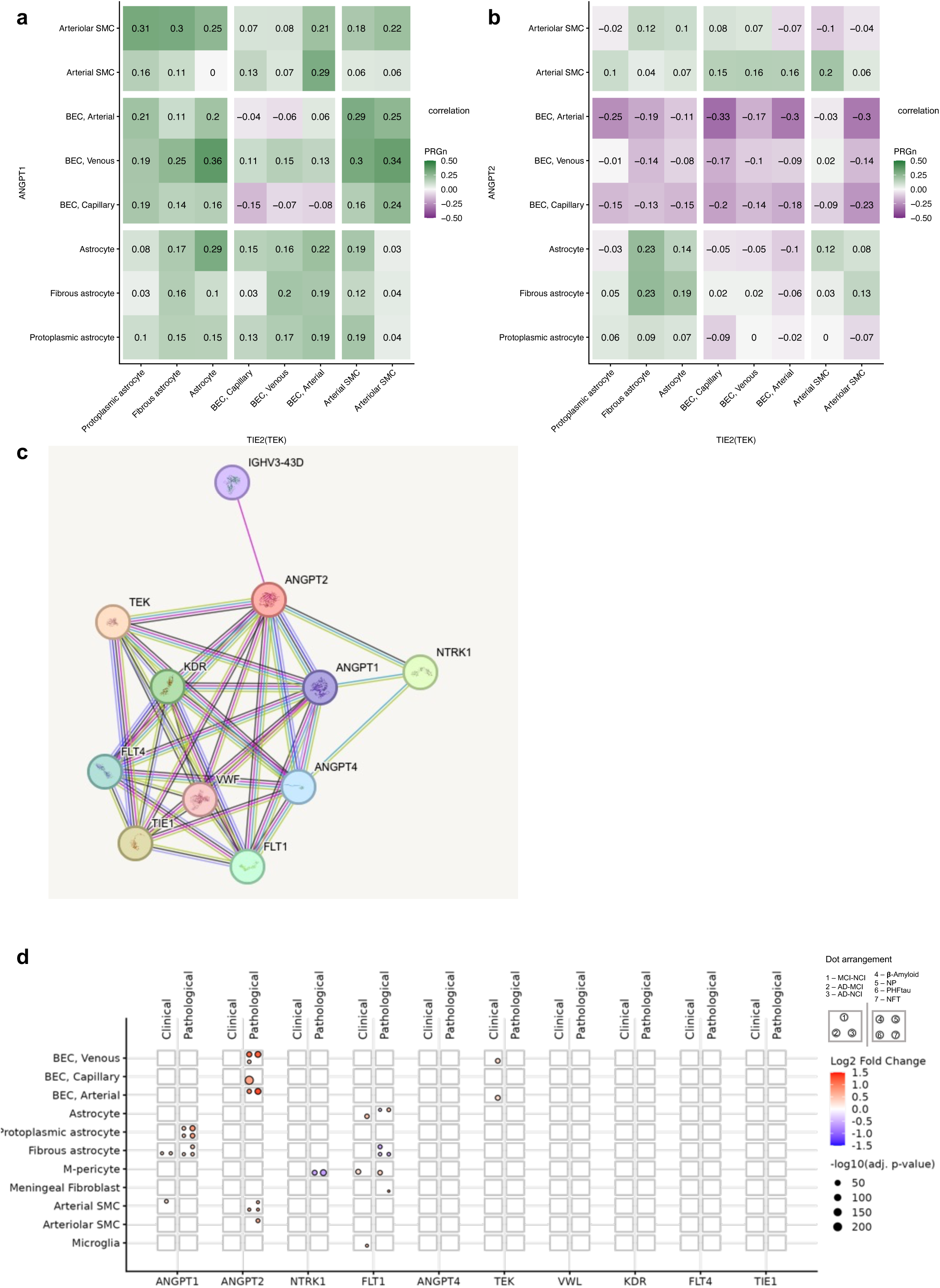
ANGPT1/2 cross cell type correlation with ANGPT1-TIE2 (**a**) and ANGPT2-TIE2 (**b**). ANGPT1 and ANGPT2 string network generated from string database (**c**). (**d**) Domino plot highlighting the string network genes present in our dataset and their dysregulation.

**Supplemental Figure 9:**
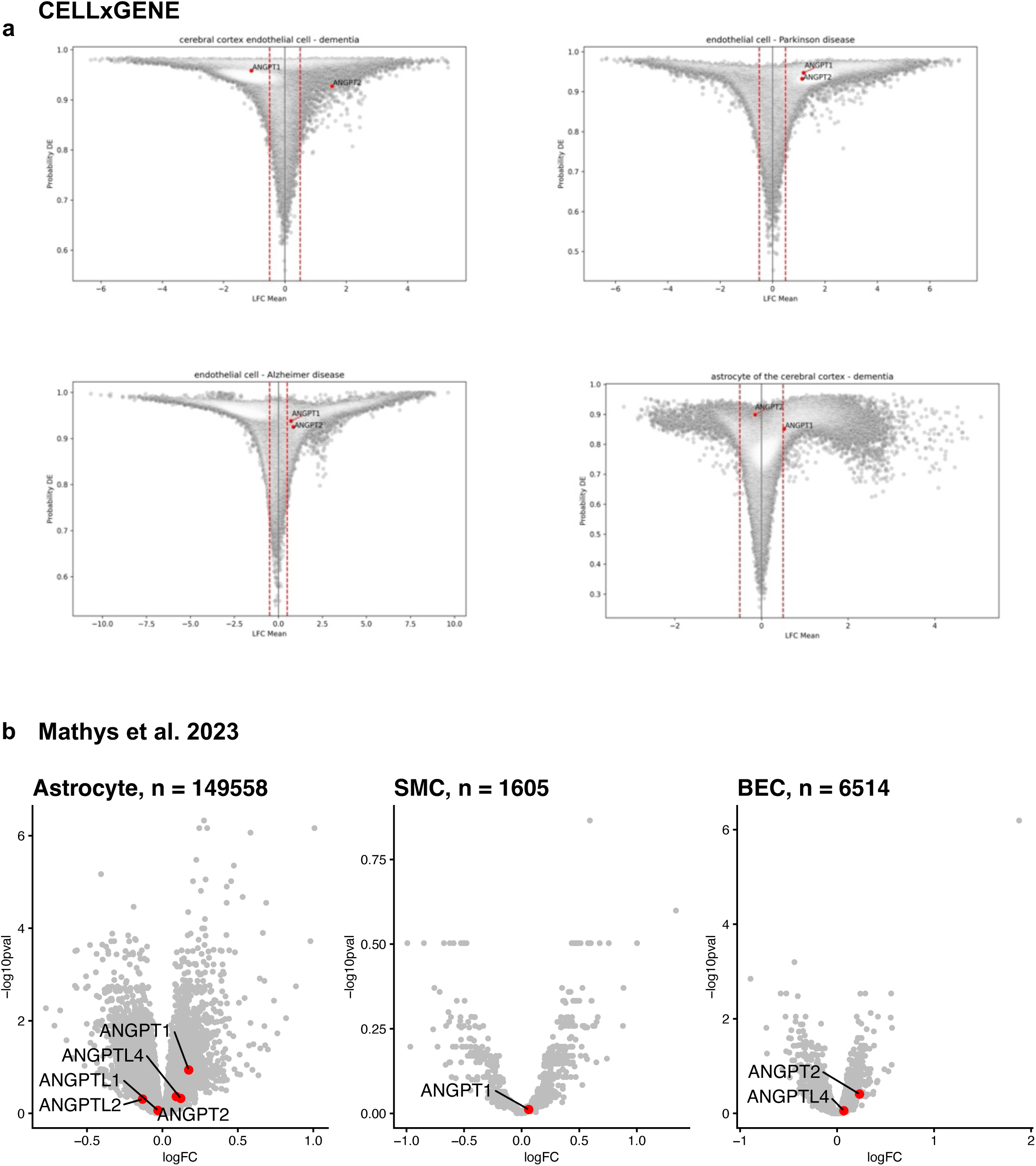
ANGPT1/2 dysregulation in other scRNA-seq studies such as the CellxGene census (**a**) and the study by Mathys et al. (**b**).

**Supplemental Figure 10:**
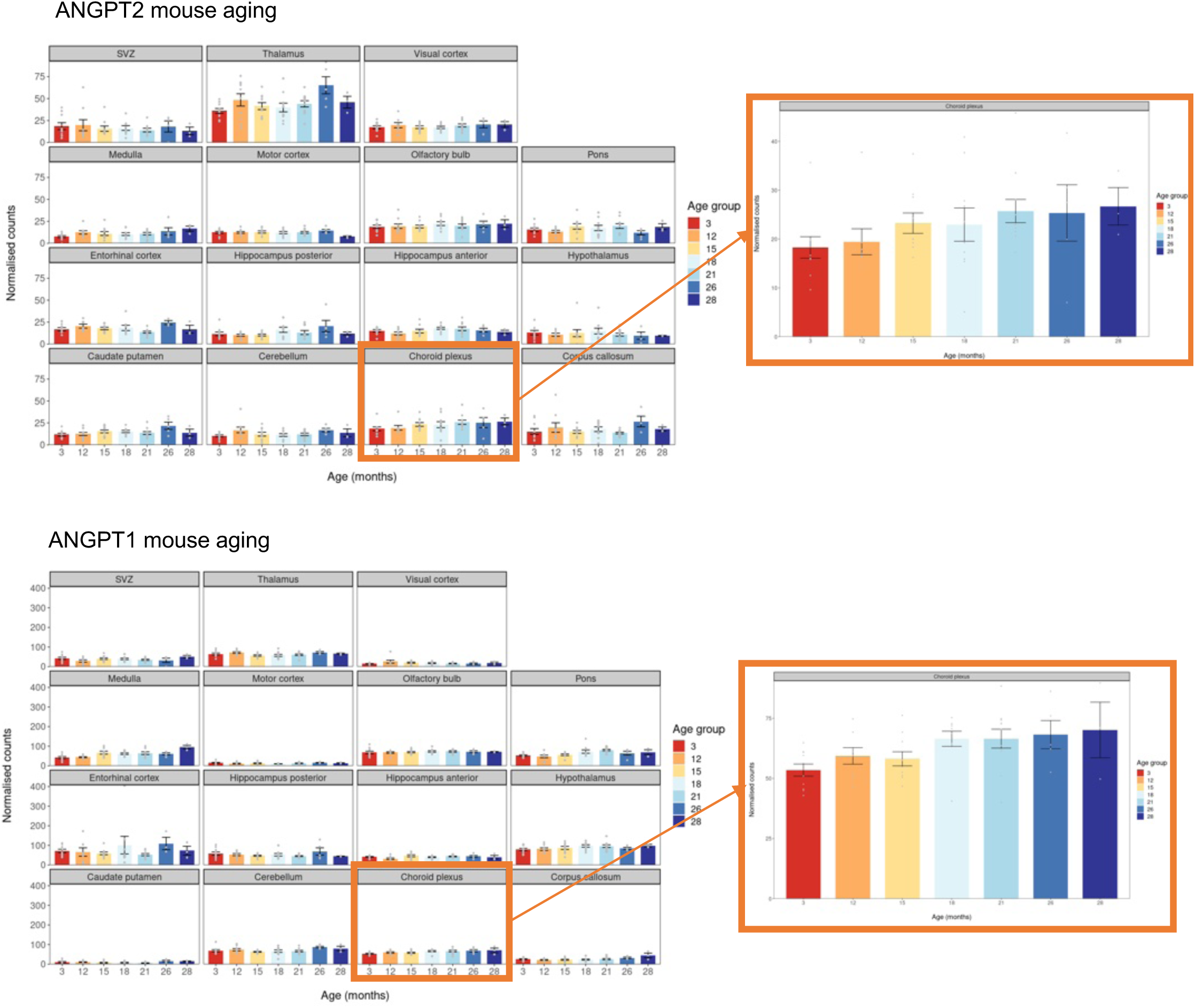
ANGPT1/2 expression mouse aging brain especially in the choroid plexus in the data published by Hahn et al.

**Supplemental Figure 11:**
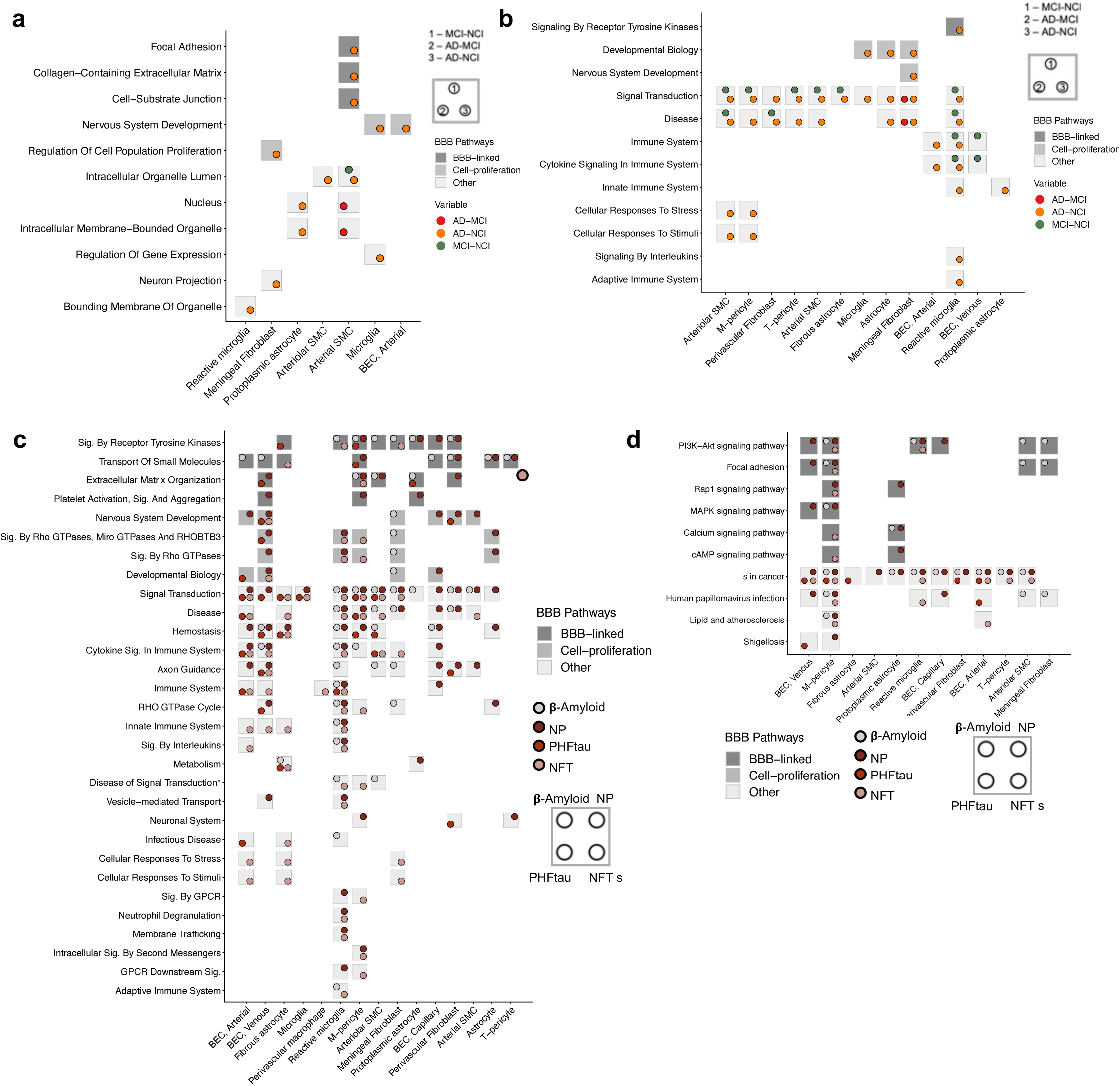
(**a,b**) Pathway DicePlots for the DEGs based on clinical diagnosis. Based on GO biological process and cellular component (**a**), Reactome 2022 (**b**), KEGG 2021 only is dysregulated in cancer signaling across all three clinical diagnosis. (**c** and **d**) Pathway DicePlots for the DEGs based on pathological plaques. Based Reactome 2022 (**c**), and KEGG 2021 (**d**).

**Supplemental Figure 12:**
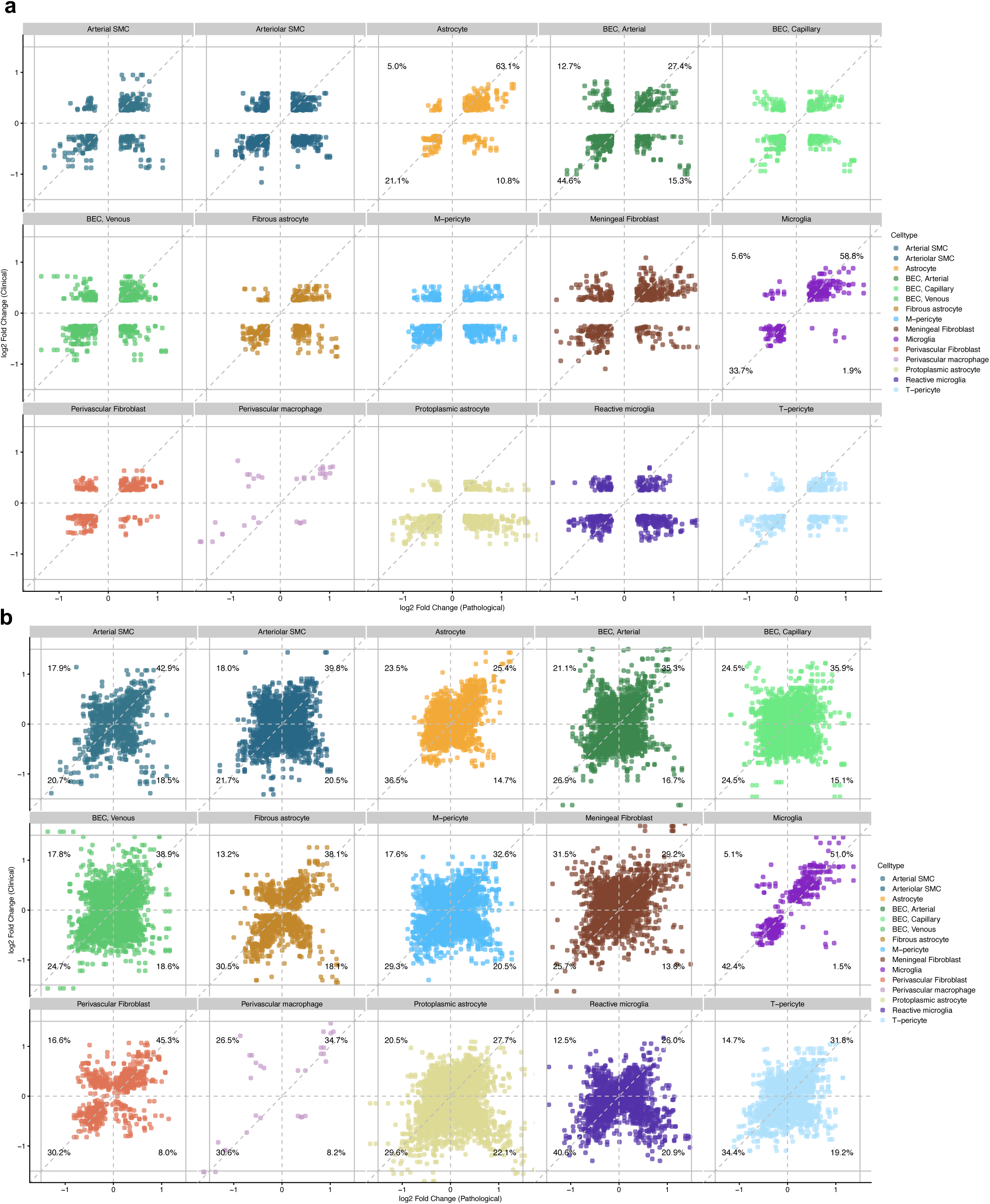
(**a**) Clinical and pathological concordance, scatterplot highlighting concordant and discordant genes per fine cell type for all significant intersecting DEGs with a logFC>0.25. (**b**) According plots as in (**a**) without a logFC threshold in the preprocessing pipeline

**Supplemental Figure 13:**
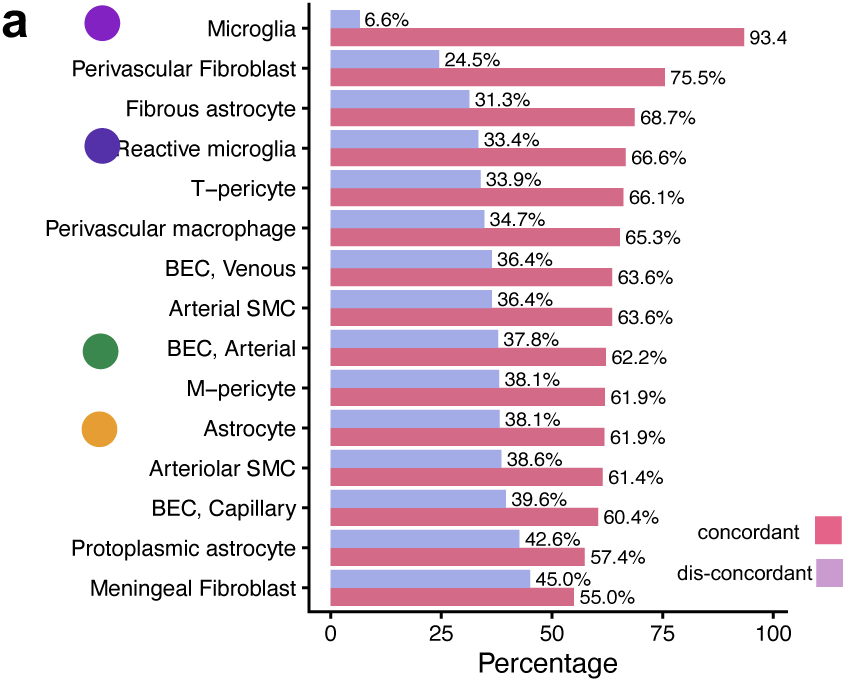
(**a**) Percentage of concordant versus discordant genes for each of the 15 cell types. The cell types are highlighted according to Figure 4 f

**Supplemental Figure 14:**
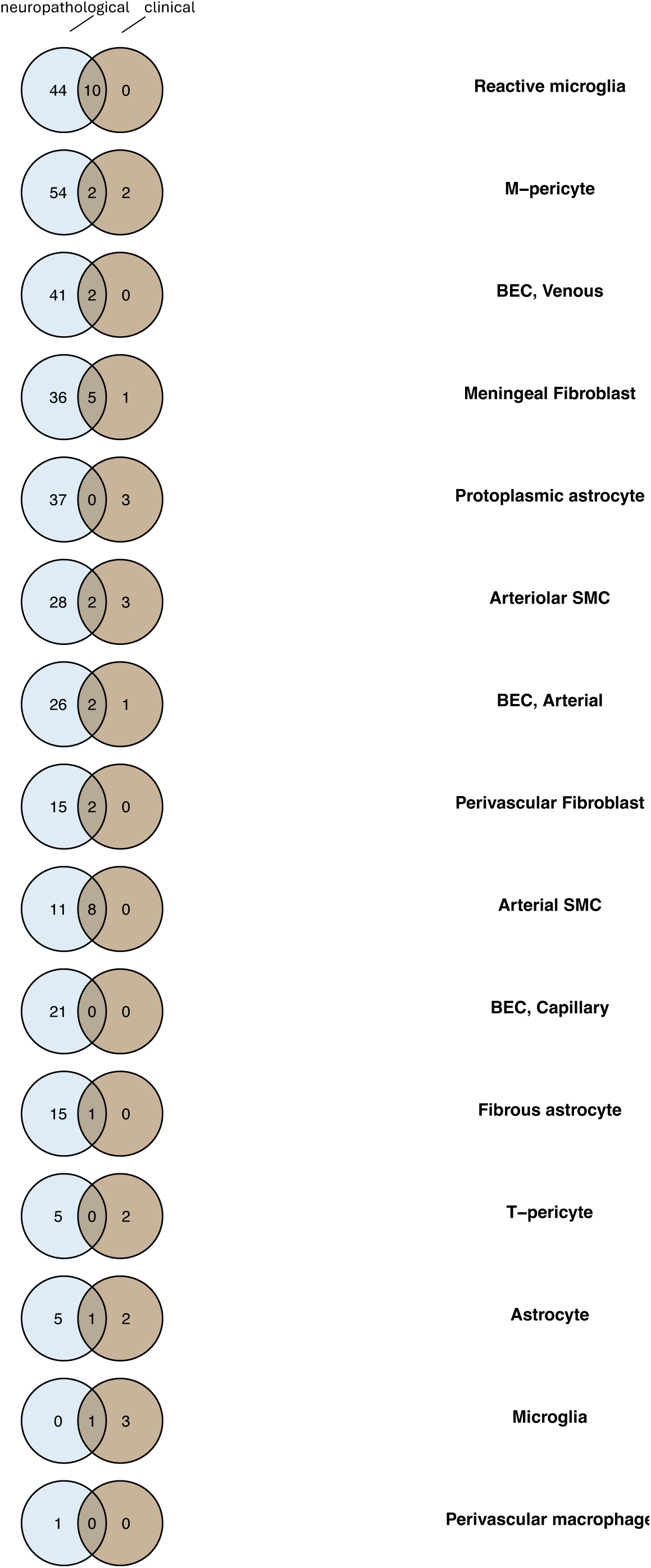
Intersection of clinical and pathological pathways, by unique pathways, i.e. pathways occurring in more than one pathological, or clinical comparison have been counted once.

**Supplemental Table 1:** Cohort metadata

**Supplemental Table 2:** NIA-Reagan DEGs filtered.

**Supplemental Table 3:** Pathology DEGs filtered.

**Supplemental Table 4**: Pathways filtered.

**Supplemental Table 5:** Clinical diagnosis DEGs.

**Supplemental Table 6:** “BBB-linked”, “Proliferation-linked”, and “Other” terms listing.

**Supplemental Table 7:** Contamination markers.

